# Aurora Kinase A proximity interactome reveals centriolar satellites as regulators of its function during primary cilium biogenesis

**DOI:** 10.1101/2020.11.05.370320

**Authors:** Melis D. Arslanhan, Navin Rauniyar, John R. Yates, Elif N. Firat-Karalar

## Abstract

Aurora kinase A (AURKA) is a conserved kinase that plays crucial roles in numerous cellular processes. Although AURKA overexpression is frequent in human cancers, its pleiotropic functions and complex spatiotemporal regulation have presented challenges in its therapeutic targeting. An essential step to overcome these challenges is the identification of the full range of AURKA regulators and substrates, which are often weak and transient. Previous proteomic studies were limited in monitoring dynamic and non-mitotic AURKA interactions. Here, we generated the first *in vivo* proximity interactome of AURKA, which consisted of over 100 proteins involving multiple biological processes and cellular compartments. Importantly, AURKA had extensive proximity interactions to centriolar satellites, key regulators of the primary cilium. Affinity pulldown and phosphoproteomics experiments confirmed this proximity relationship at the physical level. Loss-of-function experiments defined satellites as negative regulators of AURKA activity, abundance and localization in quiescent cells. Notably, loss of satellites increased AURKA activation at the basal body and resulted in defective cilium assembly and enhanced cilium disassembly. Collectively, our results provide a powerful resource for dissecting AURKA function and regulation and uncover proteostatic regulation of AURKA by centriolar satellites as a new regulatory mechanism for its non-mitotic functions.

## Introduction

Aurora kinase A (AURKA) is a member of the evolutionarily conserved Aurora serine/threonine kinase family and a pleiotropic regulator of cellular processes that are deregulated in cancer. Genetic amplification and overexpression of AURKA is prevalent in various solid and hematologic malignancies, and it correlates with aggressive cancer progression and poor survival (Nikonova et al., 2013). AURKA overexpression was shown to induce oncogenic phenotypes including aneuploidy, centrosome amplification, mitotic spindle defects and resistance to apoptosis in cultured cell lines and mammary tumors in mice (Nikonova et al., 2013, Treekitkarnmongkol et al., 2016, Wang et al., 2006, Zhang et al., 2004, Zhang et al., 2008). Due to its causal link to cancer, AURKA has been extensively investigated as a drug target for anti-cancer therapeutics. Although multiple small molecule inhibitors of AURKA activity such as alisertib (MLN8237) were shown to repress growth and progression of various cancers *in vitro* and *in vivo,* the inhibitors did not exhibit similar efficacies in cancer patients in clinical trials (Nikonova et al., 2013, Tang et al., 2017). An emerging therapeutic paradigm for overcoming the clinical limitations associated with monotherapies has been combining AURKA inhibitors with drugs targeting cancer-related AURKA functions and interactors (Chen and Lahav, 2016, Tang et al., 2017). Designing specific and effective combinatorial treatments necessitates the identification of the full extent of AURKA interactors and activities in cells.

Since its discovery as a mitotic kinase, AURKA has been extensively studied for its mitotic functions and regulation (Glover et al., 1995, Nikonova et al., 2013). AURKA interacts with and/or phosphorylates multiple proteins at the centrosomes and mitotic spindle such as PLK1, NDEL1, TACC and CDC25B and is required for centrosome maturation, mitotic entry and exit, spindle assembly and cytokinesis (Barr and Gergely, 2007, Dutertre et al., 2004, Joukov and De Nicolo, 2018, LeRoy et al., 2007, Mori et al., 2007). Spatiotemporal regulation of the pleiotropic AURKA functions in mitosis is achieved by tight control of its activity, localization and stability. While the levels and activity of AURKA are low during most of the cell cycle, they increase sharply during G2 phase and mitosis, during when they localize to the duplicated centrosomes and proximal spindle microtubules (Levinson, 2018). Its activation during mitosis is regulated by phosphorylation, ubiquitin-mediated degradation and interactions with activating proteins such as TPX2 (Bayliss et al., 2003, Levinson, 2018, Nikonova et al., 2013).

More recently, non-mitotic AURKA functions have been discovered, which include primary cilium biogenesis (Golemis et al., 2018, Plotnikova et al., 2012), stability and transcriptional activity of N-Myc (Buchel et al., 2017, Otto et al., 2009), DNA damage repair and replication fork stability (Byrum et al., 2019) and mitochondrial morphology and function (Bertolin et al., 2018, Kashatus et al., 2011) and neurite extension (Liem et al., 2009). Consistent with its diverse functions, AURKA has multiple cellular pools with differential kinase activity and substrate specificity, which are spatiotemporally regulated in dynamic cellular processes such as mitosis (Willems et al., 2018). In contrast to its extensive characterization during mitosis, little is known about how AURKA mediates its diverse non-mitotic functions and the mechanisms by which it is regulated in these contexts. This is in part due to lack of systematic studies for identification of the spatially and temporally constrained interactions that govern the non-mitotic regulation of AURKA. Addressing these questions is essential for designing combinatorial therapeutics targeting cancers linked to AURKA overexpression and to prevent the unanticipated adverse effects of AURKA inhibition (Kiseleva et al., 2019).

The first indication of the non-mitotic functions AURKA functions was its requirement for flagellar disassembly in *Chlamydomonas reinhardtii*, which was followed by its characterization during primary cilium biogenesis (DeVaul et al., 2017, Korobeynikov et al., 2017, Pan et al., 2004). Primary cilium is an immotile microtubule-based cellular protrusion that senses and transduces extracellular stimuli and thereby regulates growth, development and homeostasis (Conkar and Firat-Karalar, 2020, Mirvis et al., 2018). Accordingly, its structural and functional defects have been implicated in cancer and developmental disorders affecting multiple organ systems (Higgins et al., 2019, Reiter and Leroux, 2017). Primary cilium biogenesis is tightly coordinated with cell cycle. It assembles upon cell cycle exit (i.e. mitogen deprival and differentiation) and disassembles during mitosis. Loss of primary cilium in multiple cancers correlated with increased proliferation and disrupted signaling pathways, corroborating its functions in cell cycle regulation and cellular signaling (Jackson, 2011, Kim et al., 2011, Li et al., 2011). Given its dual functions at the primary cilium and during mitosis, AURKA is an important target for dissecting the link between cell cycle regulation and cilium biogenesis.

Upon induction of cilium disassembly by growth factor stimulation, AURKA is activated at the basal body by HEF1/NEDD1 (Pugacheva et al., 2007). Upon its activation, AURKA phosphorylates Histone Deacetylase 6 (HDAC6), which promotes deacetylation of axonemal microtubules and ciliary resorption. In addition to the well-characterized HEF1-AURKA-HDAC6 signaling axis, Ca+2/calmodulin (CaM) (Plotnikova et al., 2012), trichoplein (TCHP) (Inoko et al., 2012), nuclear distribution element (NDE)-like 1 (NDEL1) (Inaba et al., 2016), Pitchfork (Pifo) (Kinzel et al., 2010) and von Hipple-Lindau (VHL) (Hasanov et al., 2017) were identified as other AURKA interactors that are required for regulation of AURKA abundance and activation during cilium disassembly (Inaba et al., 2016, Inoko et al., 2012). In quiescent cells, AURKA is degraded and kept inactive to assemble and maintain primary cilium. Aberrant activation of AURKA in quiescent cells disrupts basal body biogenesis (i.e. recruitment of appendage proteins) and upregulates the ciliary resorption pathway, which together identifies AURKA as a negative regulator of cilium assembly (Pejskova et al., 2020). Despite its critical roles during cilium assembly and disassembly, the mechanisms by which AURKA localization, stability and abundance are regulated in quiescent cells remains poorly understood. For example, what are the mechanisms by which AURKA targeted to the basal body during cilium disassembly and inhibited to localize to the basal body during cilium assembly? What regulates the dynamic regulation of AURKA cellular abundance during these functions? Because multifaceted regulation of AURKA is achieved by its interactions with its spatially and temporally binding partners, addressing these questions requires the identification of the full range of its interacting partners, in particular the highly dynamic, transient ones.

Our current knowledge of the AURKA interactome is largely based on conventional proteomic and computational-based identification of its mitotic substrates and regulators (Tien et al., 2003, Deretic et al., 2018, Deb et al., 2020, Su et al., 2014, Kettenbach et al., 2013, Shu et al., 2015). Given the highly dynamic spatiotemporal regulation of AURKA interactome as well as its association with cytoskeletal structures (i.e. centrosomes, microtubules), these approaches were limited in probing the weak, transient and insoluble AURKA interactors such as enzyme-substrate interactions and interactions with cytoskeletal structures. Proximity-based labeling combined with mass spectrometry has emerged as a powerful approach to circumvent these inherent challenges of traditional biochemical methods. Comparative analysis of AP/MS and proximity-based labeling approaches for the same bait proteins (i.e. histones) showed that the resulting interactomes share a small number of interactors and complement each other for mapping comprehensive interaction landscapes (Lambert et al., 2014, Liu et al., 2018). Therefore, lack of the AURKA proximity interactome in mitosis and outside mitosis limits our understanding of the full extent of AURKA functions and regulation.

In this study, we applied the *in vivo* proximity-dependent biotin identification (BioID) approach to map the proximity interactome of AURKA of asynchronous cells. BioID makes use of the promiscuous *E. coli* biotin ligase BirA(R118G) (hereafter BirA*), which enables covalent biotinylation of proteins within about 10-20 nm proximity of BirA*-fusions of bait proteins (Arslanhan et al., 2020, Roux et al., 2012). The resulting interactome provides a powerful resource for dissecting the dynamic AURKA sub-complexes that enable its multifaceted regulation and diverse AURKA functions. Importantly, it revealed previously undescribed, extensive connections to centriolar satellites, which we verified by affinity pull-down and functional analysis. Our results described a proteostatic regulation of AURKA by centriolar satellites as a new regulatory mechanism for its functions during cilium assembly.

## Results

### BirA*-AURKA localizes and induces biotinylation at the centrosomes and spindle microtubules

To identify the AURKA proximity interactome, we applied BioID approach to human AURKA in human embryonic kidney (HEK293T) cells, which is a widely used cell line for proximity proteomics and thus makes benchmarking easier. As depicted in Fig. 1A, our experimental workflow consists of generation of cells stably expressing V5-BirA*-AURKA or V5-BirA* (control), induction of proximal biotinylation, pulldown of biotinylated proteins and mass spectrometry analysis. Since BirA* covalently marks all proteins proximal to the bait, 18 h biotin stimulation of asynchronous cells will reveal proximal relationships of AURKA during its entire lifetime, which will be representative of different cell cycle stages (Roux, 2013). To generate stable cells expression low levels of the fusion protein, we cloned an N-terminal fusion of AURKA into a lentiviral expression vector. Using lentiviral transduction, we generated HEK293T cells that stably express V5-BirA*-AURKA and V5-BirA*. We then assessed the expression of V5-BirA*-AURKA and biotinylation of its proximal interactors by immunostaining and immunoblotting for V5 to detect the fusion protein, streptavidin to detect biotinylated proteins and/or gamma-tubulin to mark the centrosome (Fig. 1B, 1C). V5-BirA*-AURKA localized to the centrosome in interphase and at the spindle poles and proximal spindle microtubules in mitosis (Fig. 1B). Importantly, in cells incubated with 50 μm biotin for 18 h, V5-BirA*-AURKA induced localized biotinylation at these structures (Fig.1B). Immunoblotting of lysates from cells stably expressing V5-BirA* and V5-BirA*-AURKA confirmed their expression as well as the successful streptavidin pulldown of the baits and the biotinylated proteins in their proximity (Fig. 1C). As expected, the levels of biotinylated proteins in lysates from V5-BirA*-AURKA and V5-BirA* cells treated with biotin was substantially higher relative to no biotin control. Together, these results show that V5-BirA*-AURKA is active and induces biotinylation of proteins at the centrosomes and microtubules.

**Fig. 1:**
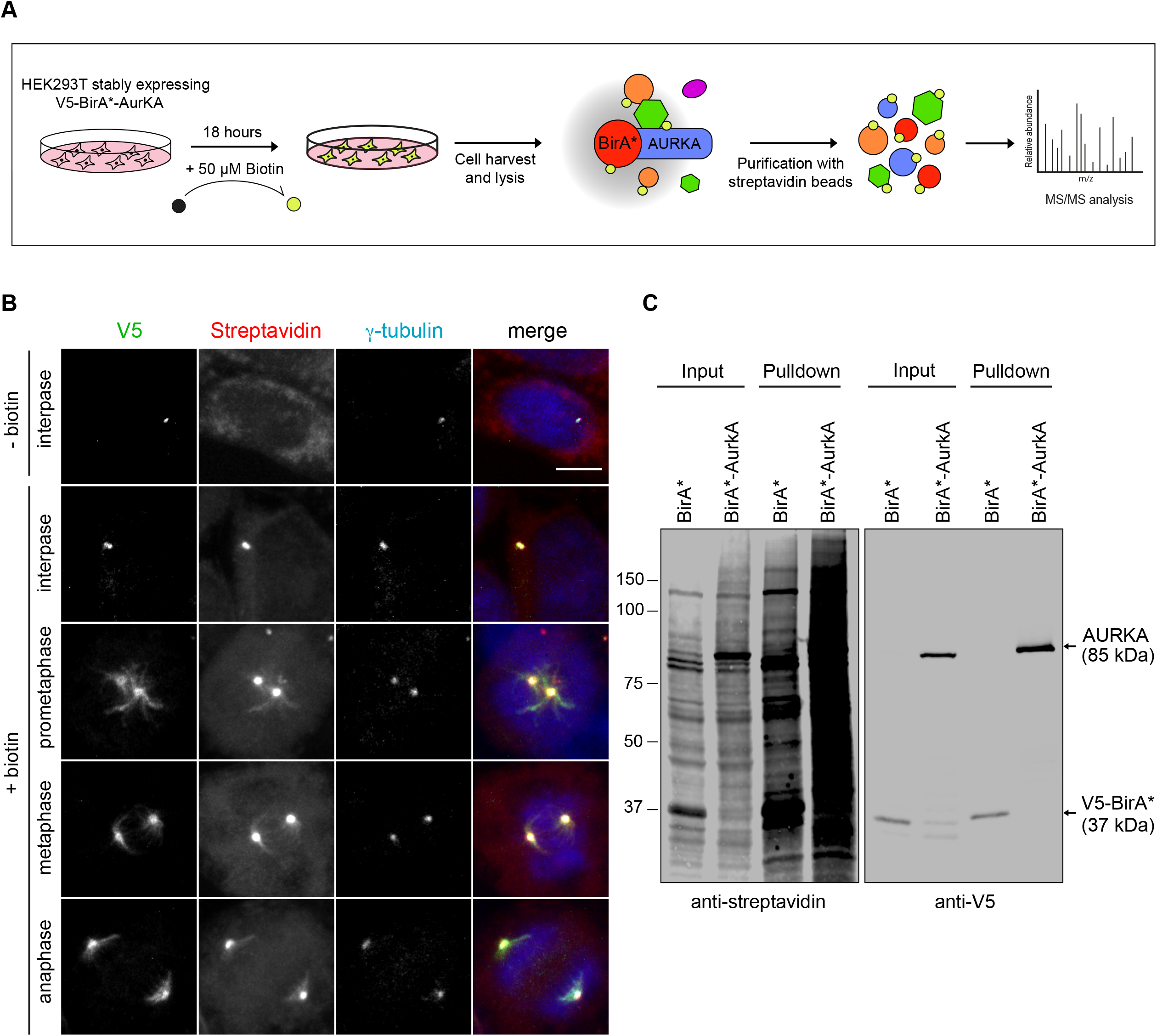
Localization and activity of BirA*-AURKA during cell cycle. **A.** Experimental workflow for the BioID experiment. HEK293T cells stably expressing V5-BirA*-AURKA and V5-BirA* were treated with 50 μM biotin for 18 hours, harvested and lysed by denaturing lysis. Biotinylated proteins were captured streptavidin affinity beads and analyzed by mass spectrometry. **B.** Biotinylation of the centrosome and microtubules by BirA*-AURKA in different cell cycle stages. After 18 h biotin incubation of HEK293T cells stably expressing BirA*-AURKA, cells were fixed and stained for the fusion protein with V5 antibody, biotinylated proteins with the fluorescent streptavidin and the centrosome with gamma-tubulin antibody. DNA was stained with DAPI. Scale bar, 10 μm **C.** Control and biotin-treated HEK293T cells stably expressing V5-BirA* and V5-BirA*-AURKA were lysed and biotinylated proteins were precipitated by streptavidin beads. The initial sample (initial) and captured biotinylated proteins (pulldown) were run on a SDS gel and immunoblotted with the fluorescent streptavidin and V5 antibody.

### Identification and validation of the AURKA proximity interactome

We performed label-free quantitative proteomics to generate the AURKA proximity interactome. For large-scale pulldowns, asynchronous cells stably expressing V5-BirA*-AURKA and V5-BirA* were grown in 5×15 cm plates and incubated with 50 μM biotin for 18 h. Following denaturing lysis of cells, biotinylated proteins were captured by streptavidin beads and analyzed by mass spectrometry. Cells expressing V5-BirA* were processed as controls. Two biological replicates for V5-BirA*-AURKA and four biological replicates for V5-BirA* were processed.

We defined the high-confidence proximity interactome of AURKA by filtering out low-confidence interactors using three different analysis methods and thresholds (Table 1). First, we performed Normalized Spectral Abundance Factor (NSAF) analysis, which accounts for run-to-run variation (Firat-Karalar et al., 2014, Zybailov et al., 2006). We included interactors with higher than 5.7 relative NSAF values of V5-BirA*-AURKA to V5-BirA* in the high confidence interactome. Second, we only accounted for proteins identified by two experimental replicates with spectral counts equal to or higher than 2. Third, we removed common mass spectrometry contaminants by using The Contaminant Repository for Affinity Purification – Mass Spectometry data (CRAPome) (Mellacheruvu et al., 2013). Altogether, these thresholds yielded 158 proteins out of the 239 proximity proteins as high confidence interactors of AURKA in asynchronous HEK293T cells.

We combined literature mining and the Biological General Repository for Interaction Datasets (BioGRID) to compile a list of AURKA interactors from low and high throughput experiments and used this list for comparative analysis with the AURKA proximity interactome (Table 2). There were 393 unique AURKA interactors deposited to BioGRID (Table 2), which were identified by affinity purification/mass spectrometry (AP/MS), phosphoproteomics, yeast two hybrid, co-immunoprecipitation (AP/immunoblotting), genetic and *in silico* experiments (Stark et al., 2006). About 10% of the high-confidence AURKA interactors (16 proteins) were reported in BioGRID, identifying the remaining 139 proteins as previously undescribed AURKA interactors (Fig. 2A, B). In addition to BioGRID analysis, we determined the AURKA proximity interactors that were previously validated as its physical interactors and/or substrates using affinity pulldown and *in vitro*/*in vivo* kinase experiments (Table 2), which identified 7 overlapping proteins (Fig. 2C). The low percentage of overlap between proximity-based proteomics and traditional approaches highlights differences in the nature of interactions probed by them and supports the use of the proximity-based labeling approach to complete the AURKA interaction landscape (Lambert et al., 2014). The AURKA interactors shared between its proximity interactome and previous studies were enriched in functions implicated in centrosome maturation, microtubule dynamics, centriole duplication, mitotic progression and cilium assembly.

**Fig. 2:**
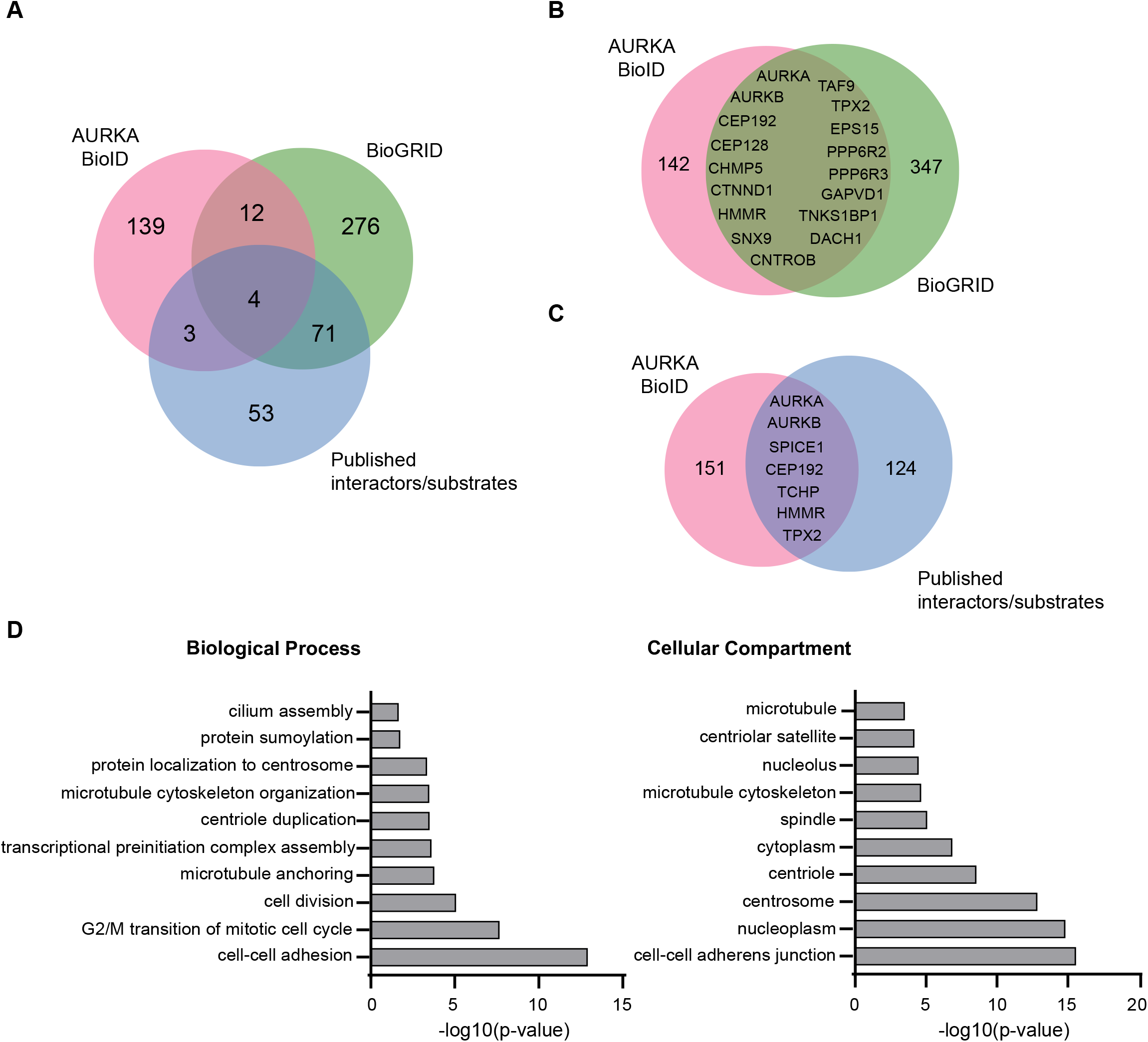
Identification and validation of high confidence AURKA proximity interactors. **A-C.** AURKA proximity interactome consists of both overlapping and distinct interactors with the published AURKA interactors. BioID-AURKA proximity interactome was compared with published physical interactors/substrates and proteins curated from the BioGRID repository. Venn diagrams show the numbers of proteins identified in the BioID-AURKA interactome color coded with pink, published interactors/substrates color coded with blue and the BioGRID interactors color coded with green. **D**. GO-enrichment analysis of the AURKA proximity interactors based on their biological process and cellular compartment. The x-axis represents the log-transformed p-value (Fisher’s exact test) of GO terms.

We performed Gene Ontology (GO) enrichment analysis of the high confidence AURKA interactors Consistent based on “biological process” and “cellular compartment” categories (Fig. 2D; Table 3). Consistent with previously described AURKA functions, GO-term analysis of biological processes showed significant (p<0.05) enrichment for cell division, microtubule-based cellular processes such as spindle assembly, DNA damage response and cilium assembly. There was also enrichment of biological processes including cell adhesion and centriole duplication, identifying them as putative new functions for AURKA. Likewise, the enriched GO-terms for cellular compartment were not limited to previously described ones such as “Centrosome” and “Spindle poles” but also included new ones such as “Centriolar Satellites” and “Cell-cell adherens junctions”. Taken together, the identification of canonical AURKA interactors (i.e. TPX2, Cep192) and GO-enrichment of AURKA-associated functions and subcellular compartments provided support for the validity of the AURKA proximity interactome and revealed a substantial number of new interactors.

### Network Analysis of AURKA interacting proteins identified by BioID

To generate a testable hypothesis for new AURKA functions and regulatory mechanisms, we organized the high-confidence AURKA proximity interactors into an interaction network by combining STRING database, ClusterONE analysis (Cytoscape) and literature mining. This analysis allowed us to group proteins based on their interconnection and thus identified the functional clusters representing subnetworks and potential multiprotein complexes. The resulting network identified a diverse array of proteins from five major functional clusters (p<0.005), which are: 1) cell cycle(30 proteins, green), 2) cytoskeleton organization (13 proteins, yellow), 3) RNA binding/processing (21 proteins, orange), 4) DNA binding (15 proteins, blue), 5) cell adhesion (14 proteins, pink) (Fig. 3A). We note that these clusters include well-characterized AURKA functions such as cell cycle as well as previously undescribed ones such as cell adhesion, and DNA binding and RNA processing/binding, linking these group of AURKA interactors to new AURKA functions.

**Fig. 3:**
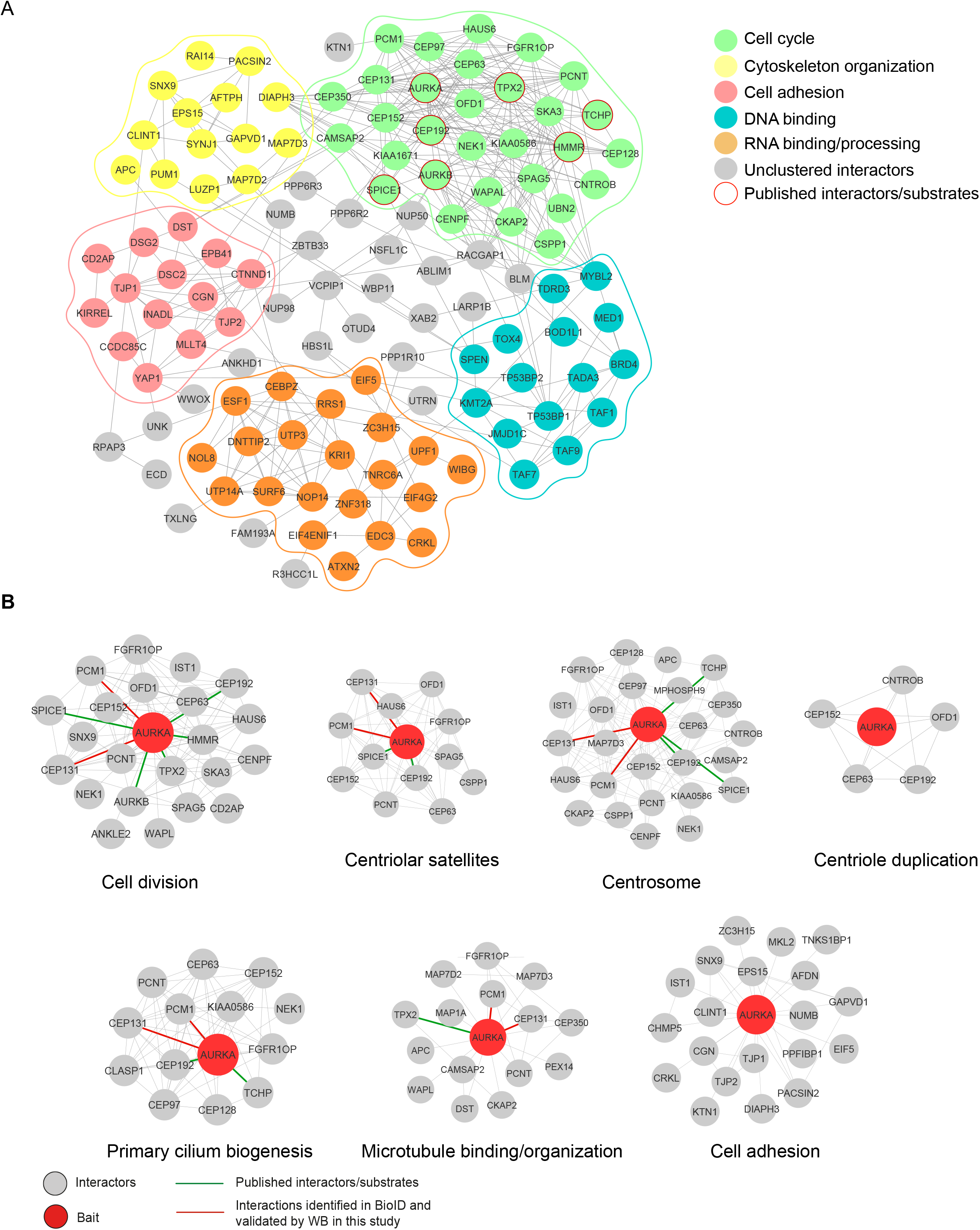
The functional interaction landscape of the AURKA proximity interactome. **A.** AURKA proximity interactome map. High confidence proximity interactors of AURKA was determined by using NSAF analysis. The interactome map was visualized in CytoScape and the functional clusters were determined by ClusterONE plug-in. Five functional clusters were enriched in the interactome, which include cell cycle (green), cytoskeleton organization (yellow), DNA binding(blue), RNA binding/processing (orange) and cell adhesion (pink). Uncategorized interactors are shown in grey and the published interactors were shown in red circles. **B.** Sub-interaction networks of AURKA based on biological process and cellular compartment. The AURKA proximity interactors were grouped using DAVID functional annotation tool and literature mining. The AURKA interactome was enriched for functions in cell cycle, centriole duplication, primary cilium biogenesis, microtubule organization and cell adhesion as well as for compartments including the centrosome and the centriolar satellites. The interconnectedness among the proteins of each network was determined by the STRING database. Nodes in green color indicates the published physical AURKA interactors/substrates and nodes in red indicates the proximity interactors validated by immunoblotting.

To account for the crosstalk between different functional clusters and to define clusters with higher resolution, we manually grouped the high confidence AURKA interactors based on their function, molecular pathway and/or cellular localization. Specifically, we generated sub-interaction interaction networks by assigning proteins based on enriched subcellular compartments (centriolar satellite and centrosome) and biological processes (cell division, centriole duplication, primary cilium biogenesis, microtubule organization, cell adhesion) (Fig. 3B). In addition to suggesting sub-complexes that work together in similar functions and localize to the same compartment, this analysis also defined the proteins that are shared between different subcellular compartments and functions. For example, a fraction of proteins that bind to microtubules function in cell division or cilium biogenesis. Moreover, there was substantial overlap between centrosome, primary cilium and centriolar satellite proteins. In contrast, proteins linked to cell adhesion did not overlap with these categories, indicative of the localization of these sub-complexes to distinct cellular compartments.

### AURKA has extensive proximity and physical interactions with centriolar satellites

AURKA proximity interactome shares a substantial number of interactors with the published proximity interactomes of the centrosome (%46) and the centriolar satellites (%38) and physical interactome of centriolar satellites (%11) (Fig. 4A, B; Table 2) (Gheiratmand et al., 2019, Gupta et al., 2015). While localization and function of AURKA at the centrosome was previously described, its nature of relationship to centriolar satellites is not known. Given that satellites interact with and regulate a wide range of centrosome and microtubule-associated proteins, we hypothesized that AURKA might be a component of centriolar satellites. To test this, we first validated the proximity relationship of AURKA to centriolar satellites by performing streptavidin pulldowns in Flag-BirA* (control) and Flag-BirA*-AURKA-expressing cells and immunoblotting with select satellite proteins. To this end, we chose satellite proteins implicated in different functions including the ciliogenesis factors PCM1, CEP131 and CEP72 and the centriole duplication protein CEP63. While PCM1, CEP131 and CEP72 co-pelleted with Flag-BirA*-AURKA, CEP63 did not (Fig 4C). This result suggests the assembly of AURKA sub-complexes with functions during ciliogenesis at the satellites. We note that the proximity interaction between PCM1 and AURKA was abolished upon inhibition of AURKA activity (Fig. S1). Finally, we tested whether the proximity interaction between AURKA and satellites are reflected at the physical level by GFP pulldown experiments from asynchronous and quiescent cells ectopically expressing GFP-PCM1 and FLAG-BirA*-AURKA (Fig. 4D). We found that endogenous FLAG-BirA*-AURKA precipitated with GFP-PCM1, but not GFP, in both asynchronous and quiescent cells (Fig 4D).

**Fig. 4:**
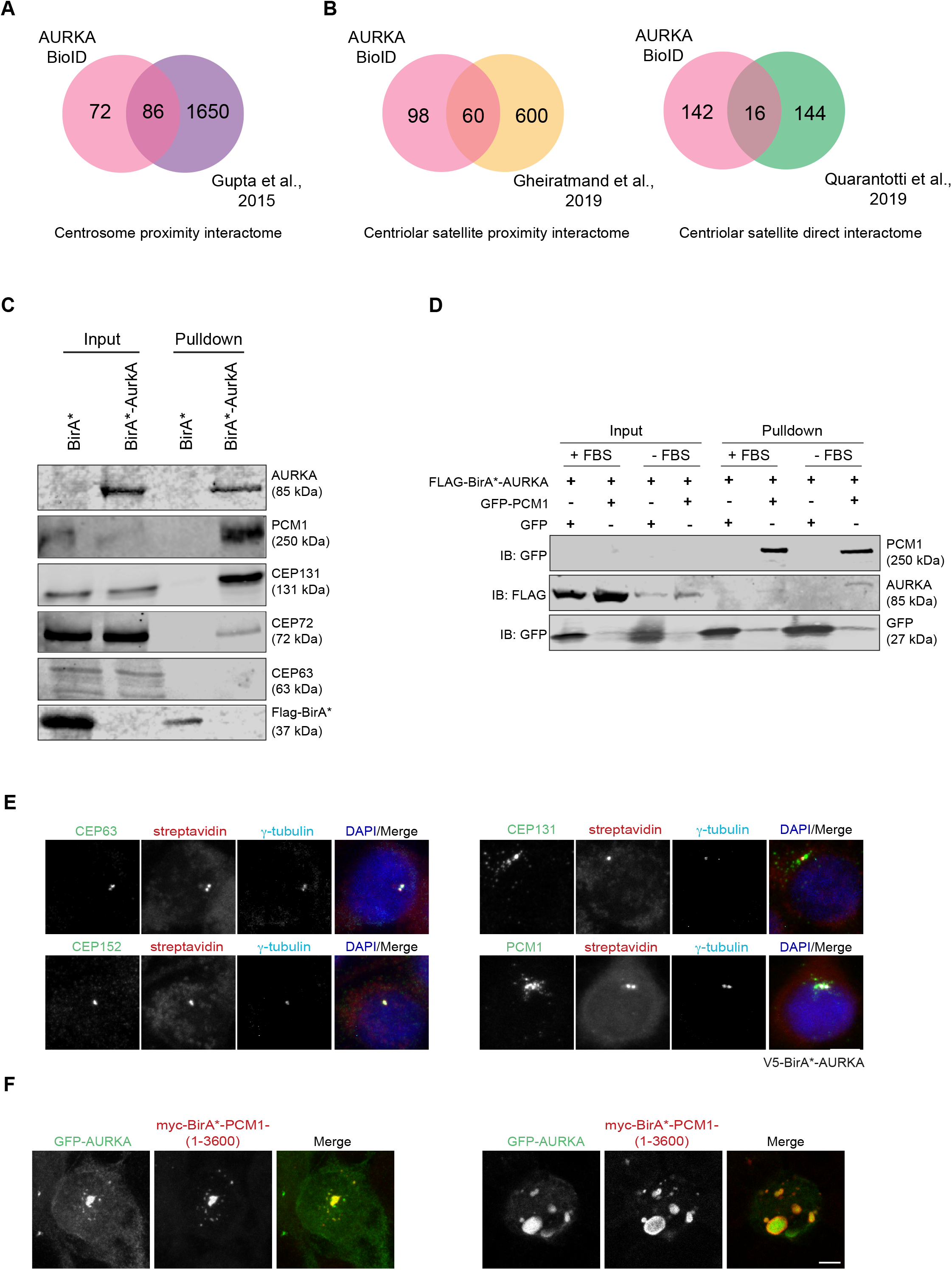
AURKA has extensive proximity interactions with centrosomes and centriolar satellites. **A-B** Comparison of the BioID-AURKA proximity interactome with the proximity interactome of the centrosome (Gupta et al., 2015), the proximity interactome of centriolar satellites (Gheiratmand et al., 2019) and the physical interactome of centriolar satellites (Quarantotti et al., 2019). **C.** AURKA interacts with the centriolar satellite proteins PCM1, CEP131 and CEP72. HEK293T cells were transiently transfected with Flag-BirA* or Flag-BirA*-AURKA. Following 18 h biotin incubation, cells were lysed, and biotinylated proteins were precipitated by streptavidin beads. The initial sample and immunoprecipitated biotinylated proteins were run on a gel and immunoblotted with fluorescent streptavidin and antibodies against FLAG, Cep72, Cep131, PCM1 and Cep63. **D.** PCM1 interacts with AURKA. HEK293T cells were transfected with FLAG-BirA*-AURKA and GFP-PCM1. Transfected cells were grown in normal or serum starved medium for 24 hours and lysed. GFP-PCM1 was immunoprecipitated with GFP-Trap beads and co-precipitated proteins were immunoblotted with antibodies against GFP and FLAG. **E.** Localization of BirA*-AURKA relative to markers of the centrosome and centriolar satellites. HEK293T cells stably expressing V5-BirA*-AURKA were incubated with biotin for 18 h and immunostained with fluorescent streptavidin and antibodies against CEP63, CEP152, CEP131 and PCM1. DNA was stained with DAPI. Scale bar, 5 μm **F.** AURKA is recruited to the cytoplasmic satellite aggregates. HEK93T cells were co-transfected with GFP-AURKA and myc-BirA*-PCM1 (1-3600). 24 h post-transfection, cells are immunostained with Myc and GFP antibodies. Scale bar, 5 μm

In parallel to interaction experiments, we determined the localization of endogenous and ectopically-expressed AURKA relative to the centrosome and cilium markers. Endogenous AURKA and V5-BirA*AURKA localized to the centrosome but not satellites, as assessed by staining for centrosome proteins CEP63 and CEP152 and satellite proteins PCM1 and CEP131 (Fig 4E). Moreover, V5-BirA*-AURKA did not induce biotinylation of satellites in biotin-treated cells. Although pull down experiments confirmed physical and proximity interactions between AURKA and satellites, localization experiments did not detect AURKA localization to satellites. This might be due to the low abundance and transient nature of this interaction. To test this, we examined whether GFP-AURKA is recruited to the cytoplasmic satellite aggregates induced by ectopic expression of PCM1(1-3600 a.a.). These aggregates were shown to disrupt satellite distribution and sequester various satellite proteins such as PLK1. We found that GFP-AURKA was recruited to the cytoplasmic satellite granules in GFP-PCM1 (1-3600)-expressing cells (Fig 4F). Taken together, interaction and localization experiments identify AURKA as a new component of satellites.

AURKA mediates its diverse functions in part through phosphorylating proteins implicated in these processes. To examine whether PCM1 is a substrate for AURKA, we expressed Flag-PCM1 in HEK293T cells and treated them with DMSO (vehicle control) or AURKA inhibitor MLN8237. Following FLAG pulldowns, in-gel digestion of FLAG-PCM1 band was performed and the resulting peptides were analyzed by mass spectrometry (Fig. 5A,B; Table 4).This analysis identified 55 phosphorylated peptides on PCM1 in control cells. Of these 55 sites, 2 of them (S988 and S1335) had the R-R-X-p[S/T] consensus sequence for AURKA and 46 were reported in previous high-throughput phosphoproteomics studies (Cheeseman et al., 2002, Hornbeck et al., 2015, Kettenbach et al., 2011). Importantly, 19 phosphopeptides were depleted in MLN8237-treated cells (Mascot score>=32, FDR<0.01), identifying these sites as putative AURKA phosphorylation sites (Fig. 5C,D,E). Together, the phosphoproteomics analysis identifies PCM1 as a putative new substrate for AURKA.

**Fig. 5:**
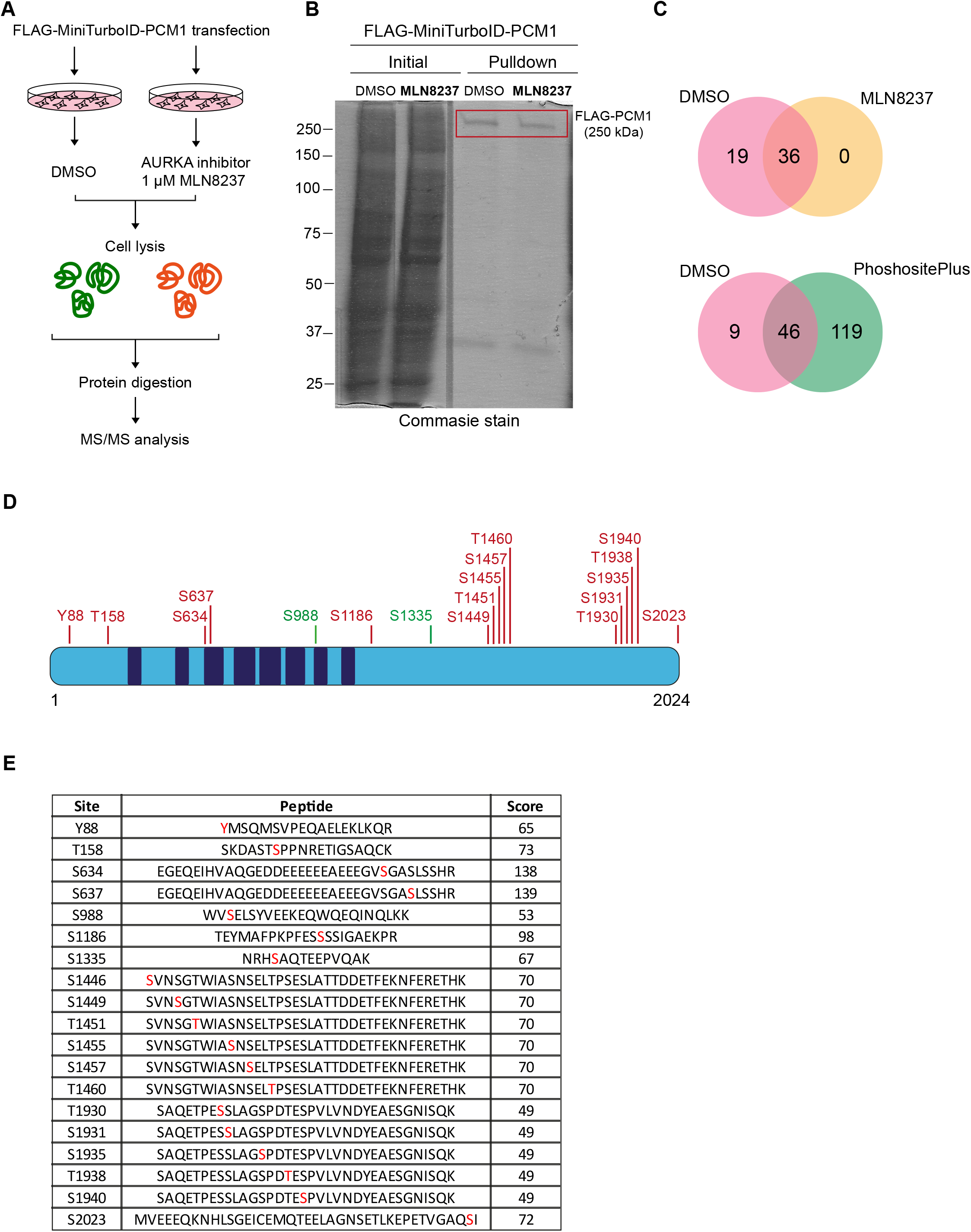
PCM1 is a putative substrate for AURKA. **A.** Experimental workflow for quantitative phosphoproteomics. HEK293T cells were transiently transfected with Flag-PCM1 (4×15 cm plates), treated either DMSO (vehicle control) or 1 μM MLN8237. Following cell lysis and FLAG pulldown, FLAG-PCM1 was excised, digested with trypsin and analyzed by mass spectrometry. **B.** Coomassie gel image of FLAG-PCM1 pulldown. The rectangle indicates the corresponding band excised for mass spectrometry. **C.** Comparison of the PCM1 phosphosites identified in DMSO (vehicle control) and MLN8237-treated cells. Venn diagrams shows the number of phosphosite. **D.** Schematic representation of phosphosites depleted in MLN8237-treated cells relative to control cells. The sites color coded with green match with the consensus AURKA motif (R/K/N-R-X-S/T). **E.** Peptides depleted in MLN8237-treated cells relative to control cells. Amino acids in red indicates the possible phosphorylation residue and the associated scores represent the Mascot peptide scores.

### Centriolar satellites regulate AURKA localization, abundance and activity during cilium biogenesis

Centriolar satellites interact with a wide range of centrosome proteins and mediate their functions at the centrosomes and primary cilium by regulating the cellular localization and abundance of their residents (Odabasi et al., 2020, Prosser and Pelletier, 2020). Given the extensive proximity relationships between satellites and AURKA and their emerging functions at the primary cilium, we hypothesized that satellites might regulate AURKA localization, stability and/or kinase activity during cilium biogenesis. To test this, we investigated how these properties are altered upon loss of satellites in human retinal pigmented epithelial (RPE1) cells, which are widely used models to study cilium biogenesis. As previously described, we induced loss of satellites by RNAi-mediated depletion of their scaffolding protein PCM1 (Kim et al., 2008).

Given that phosphorylation of AURKA on Thr288 located in its activation T loop results in its activation and increase in its enzymatic activity (Walter et al., 2000), we first examined the levels of AURKA and Thr288-phosphorylated AURKA (p-AURKA) at in control and PCM1 depleted cells serum starved for 24 h. The basal body levels of AURKA and p-AURKA increased significantly (p<0.01) in PCM1-depleted cells as compared to control cells, suggesting satellite functions in sequestration of AURKA to limit its basal body recruitment, inhibition of its activity and regulation of its cellular abundance (Fig. 6A, B). We next examined the cellular levels of AURKA by immunoblotting lysates from control and PCM1-depleted quiescent cells for AURKA and p-AURKA (Fig. 6C). Cellular AURKA and p-AURKA levels were significantly higher (p<0.01) in PCM1-depleted cells as compared with control cells, suggesting that satellites negatively regulate AURKA stability. To test this, we assayed the degradation rate of AURKA by cycloheximide block-chase experiments. While more than 53.05 ± 3.71% AURKA was degraded in control cells during 6 h cycloheximide treatment, only 10± 3.86% of PCM1 depletion was degraded (p<0.01). Inhibition of AURKA degradation suggests that satellites destabilize AURKA. Taken together, these results identify satellites as regulators of AURKA stability, activity and localization in quiescent cells.

**Fig. 6:**
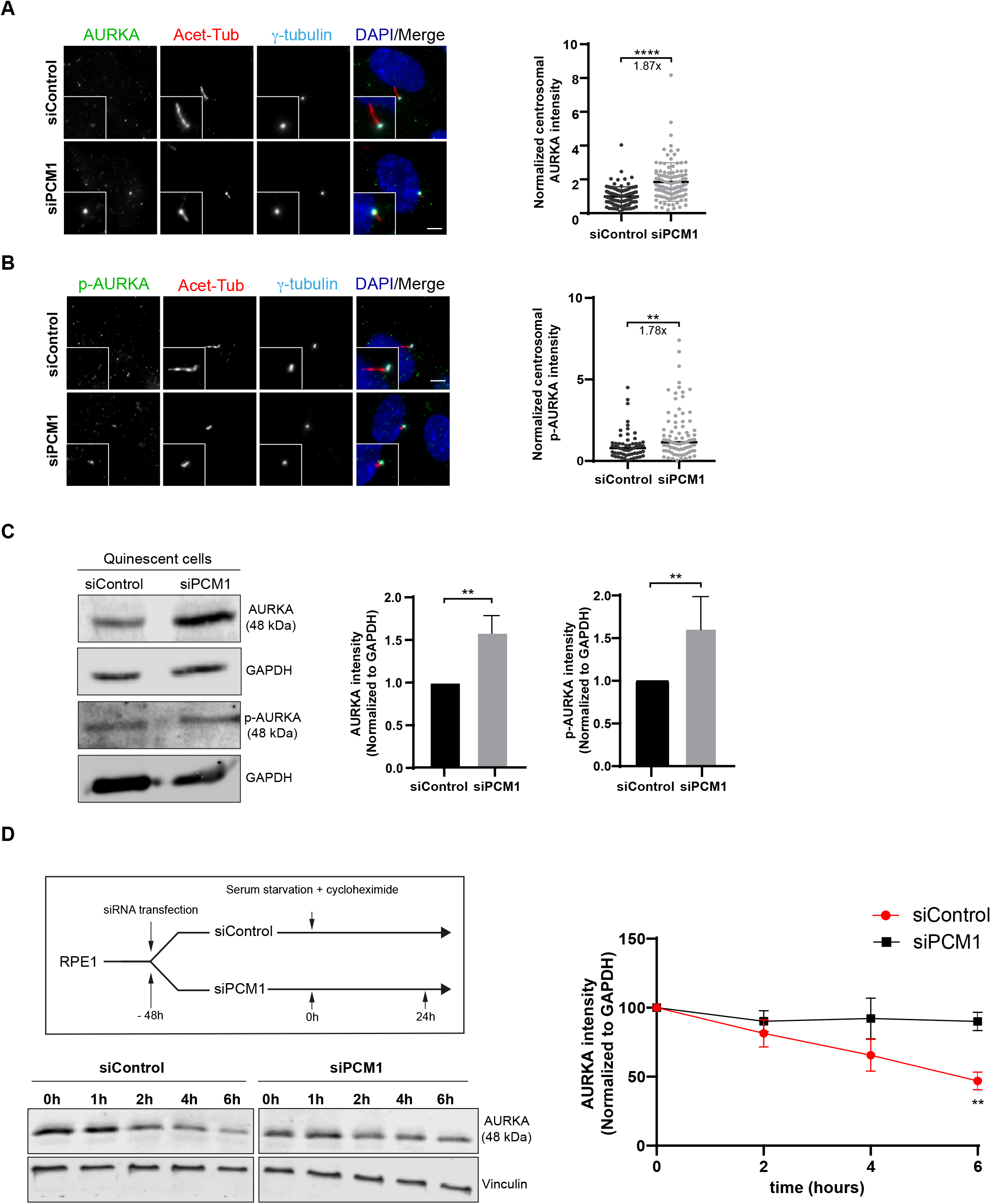
Centriolar satellites regulate AURKA localization, abundance and activity in quiescent cells. **A, B.** Representative images of basal body **(A)** AURKA and **(B)** phospho-AURKA (p-AURKA) localization in control and PCM1-depleted cells. RPE1 cells were transfected with control or PCM1 siRNA for 48 h. Following 24 h serum starvation, cells were fixed and stained with the indicated antibodies. Centrosomal AURKA and p-AURKA fluorescence intensities were measured from maximum projections, and average means of the levels in control cells were normalized to 1. n>100 cells per experiment. Data for **(A)** AURKA and **(B)** p-AURKA represent mean value from three experiments per condition ± SEM (**p < 0.01; ****p < 0.0001) **B.** Quantification of total AURKA and pAURKA levels in control and PCM1-depleted cells. RPE1 cells were transfected with control or PCM1 siRNA for 48 h. Following 24 h serum starvation, cell lysates were prepared and run out on an SDS-PAGE gel. Proteins were detected by immunoblotting with antibodies against AURKA, pAURKA and GAPDH (loading control). Data represents mean value from two experiments per condition ± SEM (**p < 0.01) **D.** Experimental workflow of experiments for assaying degradation rate of AURKA in control and PCM1-depleted cells. Cells were transfected with control or PCM1 siRNA for 48 h, then treated with 200 nM cycloheximide along with serum starvation for indicated time points. AURKA intensities were quantified by immunoblotting for AURKA and vinculin (loading control) and normalized to vinculin levels. Data represent mean value from three experiments per condition ± SEM (**p < 0.01)

### Centriolar satellites and AURKA have antagonistic functions in cilium assembly and disassembly

Having established the link between satellites and AURKA regulation in quiescent cells, we next investigated the functional consequences of this regulatory relationship. Recent studies showed that satellites function as trafficking and storage sites for key ciliogenesis factors and regulate primary cilium assembly, maintenance and disassembly (Aydin et al., 2020, Odabasi et al., 2019, Wang et al., 2016). However, the proteins regulated by satellites during cilium biogenesis have not been fully characterized. Because we showed that cellular localization, abundance and activity of AURKA is regulated by satellites, we hypothesized that satellites might mediate regulate cilium assembly and disassembly in part through AURKA. To test this, we treated control and PCM1-depleted RPE1 cells with DMSO (vehicle control) or AURKA inhibitor MLN8237 and quantified the percentage of ciliated cells 24 h post serum starvation (Fig. 7A, B, C). In agreement with previous reports, PCM1 depletion resulted in a significant decrease in the percentage of ciliated cells relative to control cells and the cilia that formed in these cells were significantly shorter (Fig. 7B,C) (siControl: 79.80 ± 6.14%, siPCM1: 31.97 ± 2.79%, p<0.001; siControl: 5.33μm ± 0.14μm; siPCM1:3.89 μm ± 0.15 μm, p<0.0001) (Aydin et al., 2020, Kim et al., 2008, Stowe et al., 2012). While MLN8237- and DMSO-treated cells ciliated with similar efficiencies (p>0.1), AURKA inhibition partially restored the ciliogenesis defect in PCM1-depleted cells (siPCM1+DMSO: 31.97 ± 2.79%; siPCM1+MLN8237: 49.70 ± 2.14%, p<0.01; siControl+DMSO: 79.80 ± 6.14%, siControl+MLN8237: 49.70 ± 7.6%, p=0.94) (Fig 7C). However, defective cilium length phenotype was not rescued by AURKA inhibition, suggesting that satellites regulate cilium length independent of AURKA activity (p=0.32) (Fig 7D). Together, these results identify AURKA activity as a mediator of cilium assembly defects in PCM1-depleted cells.

**Fig. 7:**
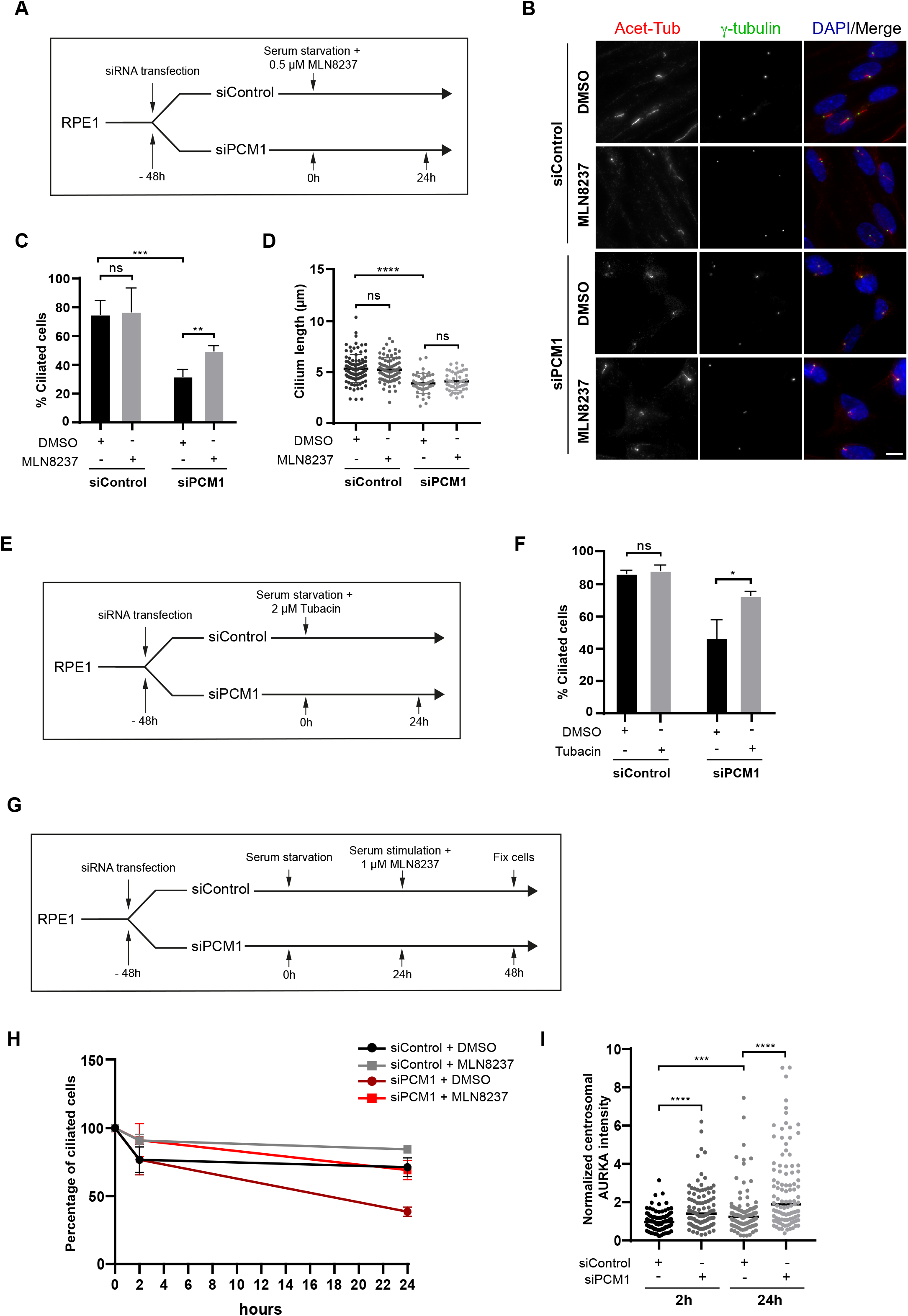
Centriolar satellites regulate cilium assembly and disassembly through inhibition of AURKA activation at the basal body. **A.** Experimental workflow of cilium assembly experiments in control and PCM1-depleted RPE1 cells treated with DMSO or MLN8237. Cells were transfected with control or PCM1 siRNA for 48 h and treated with DMSO (vehicle control) or 0.5 μM MLN8237 in serum starvation medium for 24 h. **B.** Representative immunofluorescence images for A. Cells were fixed and immunostained for the primary cilium with acetylated tubulin antibody (Acet-tub) and the centrosome with gamma-tubulin antibody. DNA was stained with DAPI. Scale bar, 10 μm **C.** Quantification of ciliogenesis efficiency for A. n>100 cells per experiment. Data represent mean value from three experiments per condition ± SEM (***p < 0.001; ns, non-significant). **D.** Quantification of cilium length for A. n>100 cells per experiment. Data represent mean value from three experiments per condition ± SEM (****p < 0.0001; ns, non-significant). **E.** Experimental workflow of cilium assembly experiments in control and PCM1-depleted RPE1 cells treated with DMSO or tubacin. Cells were transfected with control or PCM1 siRNA for 48 h and treated with DMSO (vehicle control) or 2 μM tubacin in serum starvation medium for 24 h. **F.** Quantification of ciliogenesis efficiency for A. n>100 cells per experiment. Data represent mean value from three experiments per condition ± SEM (*p < 0.05; ns, non-significant). **G.** Experimental workflow of cilium disassembly experiments in control and PCM1-depleted RPE1 cells treated with DMSO or MLN8237. Cells were transfected with control or PCM1 siRNA for 48 h, serum starved for 24 h and treated with DMSO (vehicle control) or 1 μM MLN8237 in serum stimulation medium for 2 h and 24 h. Cells were fixed and immunostained for the primary cilium and the centrosome. **H.** Quantification of cilium disassembly. *x*-axis indicates the hours after serum stimulation. n>100 cells per experiment. Data represent mean value from three experiments per condition ± SEM. Statistical analysis indicated in S3. **I.** Quantification of AURKA levels in cilium disassembly in 2 and 24 hours. n>95 cells per experiment. Data represent mean value from three experiments per condition ± SEM (***p < 0.001; ****p < 0.0001).

Activated AURKA is sufficient to induce cilium disassembly in ciliated cells (Pugacheva et al., 2007), suggesting that the ciliogenesis defect upon PCM1 depletion might be due to the induction of cilium disassembly. Given that AURKA induces cilium resorption by HDAC6 activation, we further tested this by performing rescue experiments with the HDAC inhibitor tubacin (Fig 7E). Like AURKA, inhibition of HDAC6 activity partially restored the cilium assembly defects of PCM1-depleted cells (siPCM1+DMSO: 46.87 ± 6.9%, siPCM1: 72.95 ± 1.56%, p<0.05) (Fig 7F, S2A). Of note, there was no difference in the ciliation efficiency of DMSO and tubacin-treated cells (p=0.59, siControl+DMSO: 86.63 ± 1.5μ, siPCM1: 88.47± 2.5%). These results are consistent with activation of AURKA/HDAC6 pathway and cilium disassembly pathway in PCM1 depleted cells.

In recent work, we showed that acute inhibition of satellites by their displacement to the cell periphery enhanced cilium disassembly upon serum stimulation (Aydin et al., 2020). Since this result identifies satellites as regulators of cilium disassembly, their proximity relationship to AURKA might also be relevant during serum-stimulated cilium disassembly. To investigate this relationship, we first examined the consequences of PCM1 depletion during cilium disassembly (Fig 7G). While control cells deciliated about 30% (28.7±3.4%) after 24 h serum stimulation, PCM1-depleted cells deciliated about 60% (61.4± 2%) (p<0.01). Consistent with enhanced cilium disassembly, PCM1 depletion resulted in a significant increase in the basal body levels of AURKA at 2 h and 24 h post serum stimulation (p<0.0001) (Fig 7I). Importantly, MLN8237-treatment rescued the enhanced cilium disassembly in control and PCM1-depleted cells after 24 h serum stimulation (siPCM1+DMSO: 38.60 ± 2%, siPCM1+MLN8237: 69.19 ± 4.05%, p<0.05) (Fig 7H, S2B,C,D,E). We note that MLN8237 treatment also inhibited cilium disassembly in control cells, but it had a greater effect in PCM1-depleted cells. Together, these results indicate that satellites regulate the cilium disassembly through limiting the recruitment of AURKA to the basal body.

## Discussion

In this study, we identified the first *in vivo* proximity interaction map of AURKA, which is composed of over 100 proteins associated with numerous biological processes and cellular compartments. The resulting interactome identified multiple well-described AURKA interactors and revealed previously undescribed molecular and functional relationships, highlighting the diversity of AURKA functions and complex spatiotemporal regulation. By investigating the extensive proximity relationship between AURKA and centriolar satellites, we identified AURKA as a new component of satellites and showed that satellites negatively regulate its cellular abundance, localization and activation during cilium assembly and disassembly.

Since its identification as a mitotic kinase, research on AURKA has extensively focused on its characterization during mitosis (Barr and Gergely, 2007, Nikonova et al., 2013). Multiple low and high throughput conventional approaches were used to identify mitotic AURKA interactors, which fell short in providing mechanistic insight into the emerging non-mitotic functions of AURKA such as cilium assembly and disassembly (Deretic et al., 2019, Kettenbach et al., 2011, Santamaria et al., 2011). In this study, we overcame this limitation by identifying the proximity interactors of AURKA in asynchronous cells using the BioID approach. Given that BirA* covalently marks all proteins within 10-20 nm proximity of AURKA during its entire lifetime form synthesis to the endpoint of our assay, the AURKA proximity interactome included AURKA interactors across different cell cycle stages (Roux, 2013). Therefore, it provides a powerful resource for comprehensive dissection of mitotic and non-mitotic AURKA functions and mechanisms.

The AURKA proximity interactome only partially overlapped with previously described AURKA interactors, which can be explained by two reasons. First, proximity mapping and conventional approaches complement each other by monitoring different type of interactions (Lambert et al., 2014, Liu et al., 2018). Second, majority of previous proteomic studies for AURKA specifically mapped its mitotic substrates. Yet, a number of previously characterized AURKA mitotic interactors such as TPX2 and CEP192 were among the abundant AURKA proximity interactors, confirming that information on spatially and temporally-restricted interactions during cell cycle are preserved and thus represented in its proximity map. We note that the identification of a proximity interactor relies on its spatial relationship to the bait, the time spent around the bait and its cellular abundance. For example, known AURKA interactors including INCENP, MAPRE1/2,, NDC80, NINEIN were among the low confidence AURKA interactors. Therefore, the stringent criteria we used to define the high confidence proximity interactors might have left out functionally important relationships for AURKA and thereby, these group of proteins might be taken into consideration in a targeted way in future studies.

In addition to mitotic functions, the AURKA interactome was enriched for proteins implicated in previously described non-mitotic AURKA functions such as cilium biogenesis and DNA binding. Of note, we did not find enrichment for mitochondrial proteins in the AURKA proximity interactome, which might be due to inaccessibility of these interactions by the BioID or context-dependent differences (Bertolin et al., 2018, Bertolin and Tramier, 2020). Importantly, the AURKA proximity map also identified putative new functions for AURKA in RNA binding and cell adhesion. Future studies are required to investigate the importance and relevance of these new functions to AURKA and cancer phenotypes associated with AURKA overexpression.

The AURKA proximity map revealed extensive, previously undescribed interactions with centriolar satellites. The compositional overlap between the satellite interactome and AURKA proximity map corroborates crosstalk between their function and regulation. Of note, AURKA was previously identified as a low abundance proximity interactor of PCM1 in asynchronous cells (Gupta et al., 2015). Using pulldown experiments, we validated the proximity and physical interaction between AURKA and PCM1 and showed that AURKA kinase activity is required for this interaction, which might be direct or indirect. Although AURKA was sequestered in the cytoplasmic aggregated formed by ectopic expression of PCM1(1-3600 a.a.), we have not yet observed localization of endogenous or ectopically expressed AURKA to satellites. This is mostly likely due to the transient nature of this interaction, which will result in only a low percentage of total AURKA to localize to satellites. In fact, this discrepancy between localization and interaction experiments were previously reported for a number of proteins identified in the satellite interactome such as CP110 (Spektor et al., 2007), CEP162 (Wang et al., 2013b), MDM1 (Van de Mark et al., 2015), IFT74 (Bhogaraju et al., 2013). These results highlight the power of the BioID approach in accessing interactions that cannot be identified in localization and traditional interaction mapping studies. To further our understanding of AURKA, spatial AURKA interactomes for its different sub-cellular pools and temporal AURKA interactomes in response to different stimuli should be generated. To this end, biochemical purifications of different cellular compartments can be combined with proteomics approaches that have high temporal resolution such as TurboID-based proximity labeling (Arslanhan et al., 2020, Gingras et al., 2019).

The mechanisms by which AURKA centrosomal localization and levels as well as cellular abundance are dynamically regulated in response to cell cycle cues is not known. Given the functions of satellites in protein targeting and primary cilium biogenesis, we investigated the functional relevance of the interaction between satellites and AURKA at the primary cilium. In quiescent cells, loss of satellites caused increased localization and activation of AURKA at the basal body, which were manifested as cilium assembly and disassembly phenotypes. In addition to identifying AURKA as a downstream player of PCM1 during cilium biogenesis, these results showed that satellites are required for timely targeting and activation of AURKA at the basal body by sequestering it and limiting its recruitment at the basal body. A similar relationship was previously shown between PLK1 and PCM1 (Wang et al., 2013a).

In addition to their function in protein targeting, our results define satellites as negative regulators of AURKA stability, identifying them as active scaffolds for their residents. Given that a variety of enzymes including ubiquitin ligases, deubiquitinases and kinases localize to satellites, it is likely that satellites spatially concentrate enzymes with their substrates to enhance biochemical reactions. Consistent with their proteostatic functions, we previously reported that loss of satellite rewires the global proteome (Odabasi et al., 2019). Moreover, there are two reports that showed that satellites impart their functions in part by proteostatic regulation. First, PCM1 binds to and stabilizes the ATG8 family member GABARAP by preventing its proteosomal degradation induced by the E3 ubiquitin ligase MIB1 and thereby regulates GABARAP-mediated autophagy (Joachim et al., 2017). Second, satellites promote ciliogenesis by sequestering MIB1 and regulating the centrosomal abundance of the ciliogenesis factor Talpid3 (Wang et al., 2016). Analogously, satellites might prevent basal body accumulation of AURKA during cilium assembly by modulating the activity of ubiquitin ligases or deubiquitinases that regulate AURKA stability. Additionally, future dissection of the relationship between satellites and AURKA in different contexts such as mitosis will lead to a comprehensive understanding of why AURKA localizes to satellites. These studies might reveal a broader function for satellites in spatiotemporal regulation of AURKA.

AURKA activity is tightly regulated to ensure that it functions at the right time and place. To date, various proteins have been described as activators of AURKA such as AJUBA (Hirota et al., 2003), BORA (Hutterer et al., 2006), NEDD9/HEF1 (Pugacheva et al., 2007), PIFO (Kinzel et al., 2010), NDEL1 (Inaba et al., 2016), TCHP (Inoko et al., 2012), TPX2 (Kufer et al., 2002) and calmodulin (Plotnikova et al., 2012). Increased AURKA activation at the basal body inhibited cilium assembly upon serum starvation and promoted cilium disassembly upon serum stimulation. In agreement, AURKA overexpression was shown to suppress cilium assembly by interfering with basal body maturation as well as activation of the AURKA/HDAC6-mediated cilium disassembly pathway (Inoko et al., 2012, Pejskova et al., 2020). Intriguingly, key ciliogenesis factors such as CP110 and Cep97 were among abundant AURKA proximity interactors, identifying them as candidates by which AURKA regulates basal body maturation. As for the mechanisms underlying increased AURKA activity upon loss of satellites, we envision two possibilities that are not mutually exclusive. First, it might be a consequence of increased AURKA levels at the basal body and self-activation of AURKA by phosphorylation of Thr288. Second, it might be indirect through regulation of AURKA activators by satellites. Supporting the latter mechanism is the identification of NEDD1, NDEL1, TCHP1 and calmodulin in the satellite interactome (Gheiratmand et al., 2019, Gupta et al., 2015). Future studies are required to distinguish between these possibilities.

The relationship between AURKA and centriolar satellites might be two-faceted. While our results define satellites as a regulator of AURKA activity, they also raise the possibility that PCM1 might be a novel substrate for AURKA. This is supported by our phosphoproteomic analysis of control cells and cells treated with AURKA inhibitors. Phospho-regulation of PCM1 might be required for maintaining the interaction between PCM1 and AURKA, which is supported by loss of their proximity interaction upon AURKA inhibition. Alternatively, it might be required for the integrity and/or cellular distribution of satellites, thereby for their centrosomal and ciliary functions. *In vitro* studies are required to first test whether PCM1 is an AURKA substrate and if so, to map the amino acid residues on PCM1 that are phosphorylated by AURKA. Once these sites are identified, phospho-mimetic and phospho-dead mutants of PCM1 can be used to study the functional significance of PCM1 phosphorylation by AURKA.

### Experimental Procedures

#### Cell culture and transfection

hTERT-RPE1 cells were grown in DMEM/F12 50/50 (Pan Biotech, Cat. # P04-41250) supplemented with 10% fetal bovine serum (FBS, Life Technologies, Ref. # 10270-106, Lot # 42Q5283K)) and 1% penicillin-streptomycin (P/S). HEK293T cells were grown in DMEM (Pan Biotech, Cat. # P04-03590) supplemented with 10% FBS and 1% P/S. All cells were cultured at 37 °C and 5% CO2. All cell lines were tested for mycoplasma by MycoAlert Mycoplasma Detection Kit (Lonza). RPE1 cells were transfected with the plasmids using Lipofectamine Lipofectamine for DNA transfections and Lipofectamine RNAiMax for siRNA transfections according to the manufacturer’s instructions (Thermo Fisher Scientific). The scrambled control siRNA and siRNA targeting Homo sapiens PCM1 were previously described (Conkar et al., 2019). HEK293T cells were transfected with the plasmids using 1 μg/μl polyethylenimine, MW 25 kDa (PEI, Sigma-Aldrich, St. Louis, MO). Briefly, the plasmids were diluted in Opti-MEM (Invitrogen), and incubated with PEI for 40 min at room temperature. The DNA/PEI complex was added to the cells and the culture medium was replaced with fresh medium after 6 h of incubation with the transfection mix. Lentivirus expressing V5-BirA*-AURKA or V5-BirA* were produced in HEK293T using transfer vector, packaging and envelope vectors as previously described (Gurkaslar et al., 2020). HEK293T cells were infected with the lentivirus and expression of fusion proteins in stable lines were confirmed by immunofluorescence and immunoblotting.

For cilium assembly experiments, hTERT-RPE1 cells were washed twice with PBS and incubated with DMEM/F12 supplemented with 0.5% FBS for the indicated times. For cilium disassembly experiments, ciliated RPE1 cells were incubated with DMEM/F12 50/50 supplemented with 10% FBS for the indicated times. Cells were treated with 0.5 μM MLN8237 (Selleckchem, Cat#S1133, Germany) for AURKA inhibition or 2 μM tubacin (Cayman Chemicals, Cat #13691, Ann Arbor, MI) for HDAC6 inhibition for assembly and 1 μM MLN8237 for disassembly experiments. To inhibit protein translation, cells were treated with 200 nM cycloheximide (Sigma Aldrich) for the indicated times.

### Plasmids

Full-length cDNA for *Homo sapiens* AURKA (GenBank accession no: BC002499.2) was obtained from DNASU Plasmid Repository (Seiler et al., 2014). pLVPT2-V5-BirA*-AURKA, pDEST53-AURKA and pcDNA5.1-FRT/TO-FLAG-AURKA was cloned by Gateway recombination between pDONR221-AURKA and the indicated Gateway destination plasmids. *Homo sapiens* PCM1 (GenBank accession no: NM_006197) was cloned into pCDNA5.1-FRT/TO-Flag-miniTurboID using AscI and NotI restriction sites in forward and reverse primers (gift from Bettencourt-Dias laboratory, IGC Gulbenkian). pDEST53-PCM1(1-1200) was cloned by Gateway recombination between pDONR221(1-1200) and pDEST53.

### Biotin-streptavidin affinity purification

For the BioID experiments, HEK293T stably expressing V5-BirA*-AURKA or V5-BirA*-AURKA were grown in 5×15 cm plates were grown in complete medium supplemented with 50 μM biotin for 18 h. Following biotin treatment, cells were lysed in lysis buffer (20 mM HEPES, pH 7.8, 5 mM K-acetate, 0.5 mM MgCI2, 0.5 mM DTT, protease inhibitors) and sonicated. An equal volume of 4 °C 50 mM Tris (pH 7.4) was added to the extracts and insoluble material was pelleted. Soluble materials from whole cell lysates were incubated with Streptavidin agarose beads (Thermo Scientific). Beads were collected and washed twice in wash buffer 1 (2% SDS in dH2O), once with wash buffer 2 (0.2% deoxycholate, 1% Triton X-100, 500 mM NaCI, 1 mM EDTA, and 50 mM Hepes, pH 7.5), once with wash buffer 3 (250 mM LiCI, 0.5% NP-40, 0.5% deoxycholate, 1% Triton X-100, 500 mM NaCI, 1 mM EDTA and 10 mM Tris, pH 8.1) and twice with wash buffer 4 (50 mM Tris, pH 7.4, and 50 mM NaCI). 10% of the sample was reserved for Western blot analysis and 90% of the sample to be analyzed by mass spectrometry was washed twice in 50 mM NH_4_HCO_3_. Confirmation of proximity interactors of AURKA was carried out by immunoblotting for V5 or FLAG antibodies that detect fusion protein and streptavidin that detect biotinylated proteins.

### Immunofluorescence, microscopy, and quantitation

For immunofluorescence experiments, cells were grown on coverslips and fixed in either methanol or 4% paraformaldehyde in PBS for indirect immunofluorescence. After rehydration in PBS, cells were blocked in 3% BSA (Sigma-Aldrich) in PBS + 0.1% Triton X-100. Coverslips were incubated in primary antibodies diluted in blocking solution. Anti-PCM1 antibody was generated and used for immunofluorescence as previously described (Firat-Karalar et al., 2014). Other antibodies used for immunofluorescence in this study were rabbit anti-AURKA and anti-p-AURKA (Cell signaling, #3875T) at 1:1000, rabbit anti-PCM1 (homemade) at 1:1000, mouse anti-γ-tubulin (GTU-88; Sigma-Aldrich) at 1:4000, streptavidin (Life technologies),mouse anti-acetylated tubulin (Santa Cruz Biotechnology, sc-23950) at 1:5000 and mouse anti-V5 (Invitrogen).Antibodies used for western blotting were rabbit anti-AURKA and anti-p-AURKA (Cell signaling, #3875T), rabbit anti-PCM1 (Proteintech) at 1:1000, rabbit anti-Flag (Cell signaling,#2368) at 1:1000, mouse anti-vinculin (Santa Cruz Biotechnology, sc-55465) and streptavidin (Invitrogen) at 1:10000, rabbit anti-CEP131 (Bethyl Laboratories) at 1:1000, rabbit anti-CEP72 (Bethyl) at 1:1000, Rabbit anti-CEP63 (Thermo Fisher) at 1:1000, rabbit anti-GFP (homemade) and rabbit anti-GAPDH (Cell signaling #2118). Following primary antibody incubation and 3X PBS wash, coverslips were incubated with Alexa Fluor 488-, 594-, or 680-conjugated secondary antibodies (Thermo Fisher) diluted in 1:500 in blocking solution. Biotinylated proteins were detected with streptavidin coupled to Alexa Fluor 488 or 594 (Thermo Fisher). DNA was stained with 4′,6-diamidino-2-phenylindole (DAPI; 1 μg/ml). Samples were mounted using Mowiol mounting medium containing N-propyl gallate (Sigma-Aldrich). Images were acquired with a step size of 0.3 μm in 1024×1024 format with Leica DMi8 inverted fluorescent microscope or SP8 scanning confocal microscope using the HC PL APO CS2 63x 1.4 NA oil objective.

Quantitative immunofluorescence for AURKA was performed by acquiring a z stack of control and depleted cells using identical gain and exposure settings. The z-stacks were used to assemble maximum-intensity projections. The centrosome region for each cell were defined by staining for a centrosomal marker including gamma-tubulin. The region of interest that encompassed the centrosome was defined as a circle 3 μm^2^ area centered on the centrosome in each cell. Total pixel intensity of fluorescence within the region of interest was measured using ImageJ (National Institutes of Health, Bethesda, MD) (Schneider et al., 2012). Background subtraction was performed by quantifying fluorescence intensity of a region of equal dimensions in the area proximate to the centrosome. Statistical analysis was done by normalizing these values to their mean. Primary cilium formation was assessed by counting the total number of cells, and the number of cells with primary cilia, as detected by DAPI and acetylated tubulin staining, respectively. All values were normalized relative to the mean of the control cells (= 1).

### Immunoprecipitation

HEK293T cells were co-transfected with indicated plasmids. 48 post transfection, cells were washed and lysed with lysis buffer (10 mM Tris/Cl pH 7.5, 150 mM NaCl, 0.5 mM EDTA, 0.5 % Nonidet™ P40 and protease inhibitor cocktail) for 30 min. Lysates were centrifuged at 13000 rpm for 10 min at +4°C and supernatants were transferred to a tube. 100 μl from each sample was saved as input. The rest of the supernatant was immunoprecipitated with GFP-Trap beads (ChromoTek) or Anti-FLAG M2 agarose beads (Sigma Aldrich) overnight at +4°C. After washing 3x with lysis buffer, samples were resuspended in SDS containing sample buffer and analyzed by immunoblotting.

### Cell lysis and immunoblotting

Cells were lysed in 50 mM Tris (pH 7.6), 150 mM NaCI, 1% Triton X-100 and protease inhibitors for 30 min at 4°C followed by centrifugation at 15.000 g for 15 min. The protein concentration of the resulting supernatants was determined with the Bradford solution (Bio-Rad Laboratories, CA, USA). For immunoblotting, equal quantities of cell extracts were resolved on SDS-PAGE gels, transferred onto nitrocellulose membranes, blocked with TBST in 5% milk for 1 hour at room temperature. Blots were incubated with primary antibodies diluted in 5% BSA in TBST overnight at 4°C, washed with TBST three times for 5 minutes and blotted with secondary antibodies for 1 hour at room temperature. After washing blots with TBST three times for 5 minutes, they were visualized with the LICOR Odyssey Infrared Imaging System and software at 169 mm (LI-COR Biosciences). Primary antibodies used for immunoblotting were rabbit anti-AURKA and anti-p-AURKA (Cell signaling, #3875T), rabbit anti-PCM1 (Proteintech) at 1:1000, rabbit anti-Flag (Cell signaling,#2368) at 1:1000, mouse anti-vinculin (Santa Cruz Biotechnology, sc-55465). Secondary antibodies used for western blotting experiments were IRDye680- and IRDye 800-coupled and were used at 1:15000 (LI-COR Biosciences).

### Protein identification by Mass spectrometry

Proteins bound to beads were reduced by 20-minute incubation with 5 mM TCEP (tris(2-carboxyethyl) phosphine) and alkylated in the dark by treatment with 10mM Iodoacetamide for 20 additional minutes. The proteins were subsequently digested by adding Sequencing Grade Modified Trypsin (Promega, Madison, WI, USA) and placing the reaction mixture in a Thermomixer (Eppendorf, Westbury, NY) and incubating overnight at 37 °C at 750 rpm. The next day, the sample was acidified with formic acid to a final concentration of 5% and spun at 14,000 rpm for 30 min. The supernatant was carefully transferred to a separate microfuge tube so as not to disturb the bead pellet, and pressure-loaded into a biphasic trap column. MS analysis of the samples was performed using MudPIT technology [S7]. Capillary columns were prepared in-house from particle slurries in methanol. An analytical column was generated by pulling a 100 μm ID/360 μm OD capillary (Polymicro Technologies, Inc., Phoenix, AZ) to 3 μm ID tip. The pulled column was packed with reverse phase particles (Aqua C18, 3 μm dia., 90 Å pores, Phenomenex, Torrance, CA) until 15 cm long. A biphasic trapping column was prepared by creating a Kasil frit at one end of an undeactivated 250 μm ID/360 μm OD capillary (Agilent Technologies, Inc., Santa Clara, CA), which was then successively packed with 2.5 cm strong cation exchange particles (Partisphere SCX, 5 μm dia., 100 Å pores, Phenomenex, Torrance, CA) and 2.5 cm reverse phase particles (Aqua C18, 5 μm dia., 90 Å pores, Phenomenex, Torrance, CA). The trapping column was equilibrated using buffer A (5% acetonitrile/0.1% formic acid) prior to sample loading. After sample loading and prior to MS analysis, the resin-bound peptides were desalted with buffer A by letting it flow through the trap column. The trap and analytical columns were assembled using a zero-dead volume union (Upchurch Scientific, Oak Harbor, WA). LC−MS/MS analysis was performed on LTQ Orbitrap or LTQ Orbitrap Velos (Thermo Scientific, San Jose, CA, USA) interfaced at the front end with a quaternary HP 1100 series HPLC pump (Agilent Technology, Santa Clara, CA, USA) using an in-house built electrospray stage. Electrospray was performed directly from the analytical column by applying the ESI voltage at a tee (150 μm ID, Upchurch Scientific) directly downstream of a 1:1000 split flow used to reduce the flow rate to 250 nL/min through the columns. A fully automated 6-step MudPIT run was performed on each sample using a three mobile phase system consisting of buffer A (5% acetonitrile/0.1% formic acid), buffer B (80% acetonitrile/0.1% formic acid), and buffer C (500 mM ammonium acetate/5% acetonitrile/0.1% formic acid). The first step was 60 min reverse-phase run, whereas five subsequent steps were of 120 min duration with different concentration of buffer C run for 4 min at the beginning of each of the gradient. In LTQ Orbitrap Velos, peptides were analyzed using a Top-20 data-dependent acquisition method in which fragmentation spectra are acquired for the top 20 peptide ions above a predetermined signal threshold. As peptides were eluted from the microcapillary column, they were electrosprayed directly into the mass spectrometer with the application of a distal 2.4 kV spray voltage. For each cycle, full−scan MS spectra (m/z range 300-1600) were acquired in the Orbitrap with the resolution set to a value of 60,000 at m/z 400 and an automatic gain control (AGC) target of 1×106 ions and the maximal injection time of 250 ms. For MS/MS scans the target value was 10,000 ions with injection time of 25 ms. Once analyzed, the selected peptide ions were dynamically excluded from further analysis for 120 s to allow for the selection of lower-abundance ions for subsequent fragmentation and detection using the setting for repeat count = 1, repeat duration = 30 ms and exclusion list size = 500. Charge state filtering, where ions with singly or unassigned charge states were rejected from fragmentation was enabled. The minimum MS signal for triggering MS/MS was set to 500 and an activation time of 10 ms were used. All tandem mass spectra were collected using normalized collision energy of 35%, an isolation window of 2 Th. In LTQ Orbitrap, peptides were analyzed using a Top-10 data-dependent acquisition method. For protein identification we used Integrated Proteomics Pipeline (IP2, San Diego, CA) software, a web-based proteomics data analysis platform that supports both cloud and cluster computing, developed by Integrated Proteomics Applications, Inc. (http://www.integratedproteomics.com/). Tandem mass spectra were extracted from the Xcalibur data system format (.raw) into MS2 format using RawXtract1.9.9.2. The MS/MS spectra were searched with the ProLuCID algorithm against the EBI human IPI database (version 3.71, release date March 24, 2010) that was concatenated to a decoy database in which the sequence for each entry in the original database was reversed. The database also had sequence for two proteins, *E. coli* BirA-R118G and mouse CCDC67, appended to it. The search parameters include 50 ppm peptide precursor mass tolerance and 0.6 Da for the fragment mass tolerance acquired in the ion trap. The initial wide precursor mass tolerance in the database search was subjected to post search filtering and eventually constrained to 20 ppm. Carbamidomethylating on cysteine was defined as fixed modification and phosphorylation on STY was included as variable modification in the search criteria. The search space also included all fully− and semi−tryptic peptide candidates of length of at least six amino acids. Maximum number of internal miscleavages was kept unlimited, thereby allowing all cleavage points for consideration. ProLuCID outputs were assembled and filtered using the DTASelect2.0 program that groups related spectra by protein and removes those that do not pass basic data-quality criteria [S8]. DTASelect2.0 combines XCorr and ΔCN measurements using a quadratic discriminant function to compute a confidence score to achieve a user-specified false discovery rate (less than 1% in this analysis).

### Mass Spectrometry data analysis for BioID experiments

Raw data of two biological replicates for V5-BirA*-AURKA and four biological replicates for V5-BirA* were analyzed by filtering non-specific proteins identified from experimental control sets as well as common MS background proteins. To compare data across different runs and calculate the relative abundance of each proximity partner, we used Normalized Spectral Abundance Factor (NSAF) analysis as previously described (Firat-Karalar et al., 2014). The spectral counts of each proteins were divided to the sum of spectral counts of all proteins resulting the normalized spectral counts. We did not take into account protein length for calculating normalized values for each protein. To distinguish the nonspecific interactors from the specific ones, we calculated the ratio of normalized values of each protein in V5-BirA*-AURKA dataset relative to its normalized value in V5-BirA* dataset. A protein was considered a contaminant if the ratio was smaller than 1. Following NSAF analysis, we removed proteins identified in only one experiment or with spectral counts less than 2. Finally, we used The Contaminant Repository for Affinity Purification – Mass Spectometry data (CRAPome) to eliminate the common mass spectrometry contaminants. As the CRAPome cut-off, we removed proteins identified in >45 experiments out of 411 as contaminants. Finally, we defined the high confidence AURKA interactome as the proteins that have NSAF values in the V5-BirA*-AURKA interactome greater than 5.7 relative to the V5-BirA* interactome. We determined this cut-off value empirically by confirming the inclusion of the validated AURKA interactors in the high confidence list.

Comparative analysis with BioID results and literature were carried out incorporating interactors in BioGRID database (whole proteome analysis) as well as experimentally published physical interactors/substrates (data from *in vitro* analyses, co-immunoprecipitation etc.).

### Network modelling and clustering analysis

NSAF analysis returned 158 high confidence interactors of AURKA. We performed the network analyses for thehigh confidence interactors using the STRING database and literature mining. The network output file was visualized using Cytoscape 3.7.2. Cluster analysis were carried out by Clustering with Overlapping Neighborhood Expansion (ClusterONE) plug-in of Cytoscape and proteins were grouped together based on their function or associated cellular compartment to reveal overlapping complexes (Nepusz et al., 2012). GO terms were determined by using Database for Annotation, Visualization and Integrated Discovery (DAVID).

### Identification of phosphorylated peptides

Coomassie-stained bands from two experimental replicates from FLAG-PCM1 pulldown experiments were excised, chopped into small pieces and transferred to 1.5 ml Eppendorf tubes. For all following steps, buffers were exchanged by two consecutive 15 min incubation steps of the gel pieces with 200 μl of acetonitrile (ACN) whereby the ACN was removed after each step. Proteins were reduced by the addition of 200 μl of a 10 mM DTT solution in 100 mM ammonium bicarbonate (AmBiC, Sigma Aldrich, A6141) and incubation at 56°C for 30 min. Proteins were alkylated by the addition of 200 μl of a 55 mM chloroacetamide (CAA) in 100 mM AmBiC and incubation for 20 min in the dark. A 0.1 μg/μl stock solution of trypsin (Promega, V511A) in trypsin resuspension buffer (Promega, V542A) was diluted with ice-cold 50 mM AmBiC buffer to achieve a final concentration of 1 ng/μl. 50 μl thereof were added to gel pieces, which were incubated first for 30 min on ice and then over night at 37°C. Gel pieces were sonicated for 15 min, spun down and the supernatant was transferred into a glass vial (VDS Optilab, 93908556). Remaining gel pieces were washed with 50 μl of an aqueous solution of 50% ACN and 1% formic acid and sonicated for 15 min. The combined supernatants were dried in a speedvac and reconstituted in 10 μl of an aqueous solution of 0.1% (v/v) formic acid.

Peptides were analyzed by LC-MS/MS on an Orbitrap Fusion Lumos mass spectrometer (Thermo Scentific) as previously described (PMID:30858367). To this end, peptides were separated using an Ultimate 3000 nano RSLC system (Dionex) equipped with a trapping cartridge (Precolumn C18 PepMap100, 5 mm, 300 μm i.d., 5 μm, 100 Å) and an analytical column (Acclaim PepMap 100. 75 × 50 cm C18, 3 mm, 100 Å) connected to a nanospray-Flex ion source. The peptides were loaded onto the trap column at 30 μl per min using solvent A (0.1% formic acid) and peptides were eluted using a gradient from 2 to 85% Solvent B (0.1% formic acid in acetonitrile) over 30 min at 0.3 μl per min (all solvents were of LC-MS grade). For the detection of posttranslational modifications peptides were eluted using a gradient from 2 to 80% solvent B over 60 min. The Orbitrap Fusion Lumos was operated in positive ion mode with a spray voltage of 2.2 kV and capillary temperature of 275 °C. Full scan MS spectra with a mass range of 375–1200 m/z were acquired in profile mode using a resolution of 120,000 (maximum injections time of 50 ms, AGC target was set to 400% and a max injection time of 86 ms. Precursors were isolated using the quadrupole with a window of 1.2 m/z and fragmentation was triggered by HCD in fixed collision energy mode with fixed collision energy of 34%. MS2 spectra were acquired with the Orbitrap with a resolution of 30.000 and a max injection time of 86 ms.

The Orbitrap Fusion Lumos was operated in positive ion mode with a spray voltage of 2.2 kV and capillary temperature of 275 °C. Full scan MS spectra with a mass range of 350–1500 m/z were acquired in profile mode using a resolution of 120,000 (maximum injections time of 100 ms, AGC target was set to standard and a RF lens setting of 30%. Precursors were isolated using the quadrupole with a window of 1.2 m/z and Fragmentation was triggered by HCD in fixed collision energy mode with fixed collision energy of 30%. MS2 spectra were acquired in Ion Trap normal mode. The dynamic exclusion was set to 5 s. Acquired data were analyzed using IsobarQuant (PMID: 26379230) and Mascot V2.4 (Matrix Science) using a reversed Uniprot homo sapiens database (UP000005640) including common contaminants.

### Statistical analysis

Statistical results, average and standard deviation values were computed and plotted by using Prism (GraphPad, La Jolla, CA). Two-tailed t-tests, one-way and two-way ANOVA tests were applied to compare the statistical significance of the measurements. Error bars reflect SD. Following key is followed for asterisk placeholders for *p*-values in the figures: \**P* < 0.05, \*\**P* < 0.01, \*\*\**P* < 0.001, **** *P* < 0.0001

## Acknowledgements

We acknowledge the Firat-Karalar lab members for insightful discussions regarding this work. We acknowledge Ahmet Aktas for cloning the pCDNA5.1-FRT/TO-FLAG-miniTurboID-PCM1 plasmid, Monica Bettencourt-Dias (IGC Gulbenkian) for sharing the pCDNA5.1-FRT/TO-Flag-miniTurbo plasmid and Sebastian Patzke (University of Oslo) for RPE1-hTERT cells. We acknowledge the European Molecular Biology Laboratory (Heidelberg) for the use of the Proteomics Core Facility, which performed the identification of phosphorylated peptides. This work was supported by European Research Council (ERC) grant 679140 ENF, European Molecular Biology Organization (EMBO) Installation grant 3622 and an EMBO Young Investigator Award to ENF and the National Institutes of Health Grant to JRY.

## Author Contributions

Conceptualization, ENF, MDA.; Methodology, ENF, MDA, NR.; Investigation (light microscopy, molecular biology, and biochemical analysis), MDA.; Investigation (proximity labeling and mass spectrometry), ENF, NR.; Investigation (network analysis), MDA.; Resources, ENF, MDA.; Writing—Original Draft, ENF, MDA.; Writing—Review & Editing, ENF, MDA.; Visualization, MDA.; Supervision, ENF.; Project Administration, EBF.; Funding Acquisition, ENF, JRY.

---

**Supp. Fig. 1:**
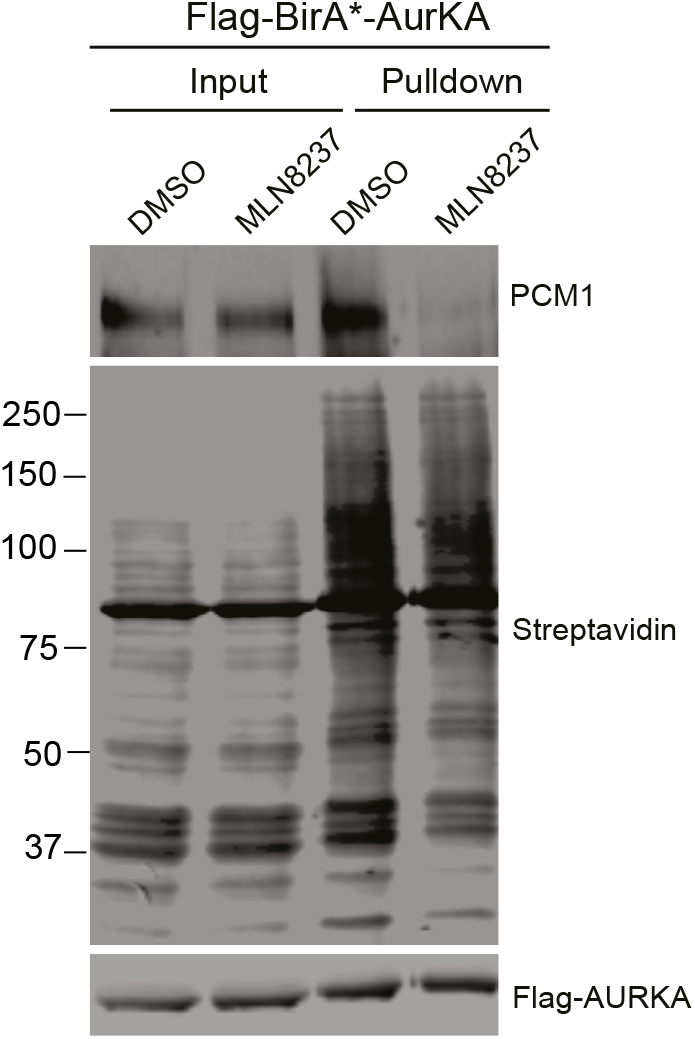
AURKA and PCM1 interaction requires AURKA activity. HEK293T cells were transiently transfected with Flag-BirA*-AURKA. 24 h post transfection, cells were treated with DMSO+biotin or MLN8237+biotin, lysed and biotinylated proteins were precipitated by streptavidin beads. The initial sample and immunoprecipitated biotinylated proteins were run on a gel and immunoblotted with fluorescent streptavidin and antibodies against FLAG and PCM1.

**Supp. Fig. 2:**
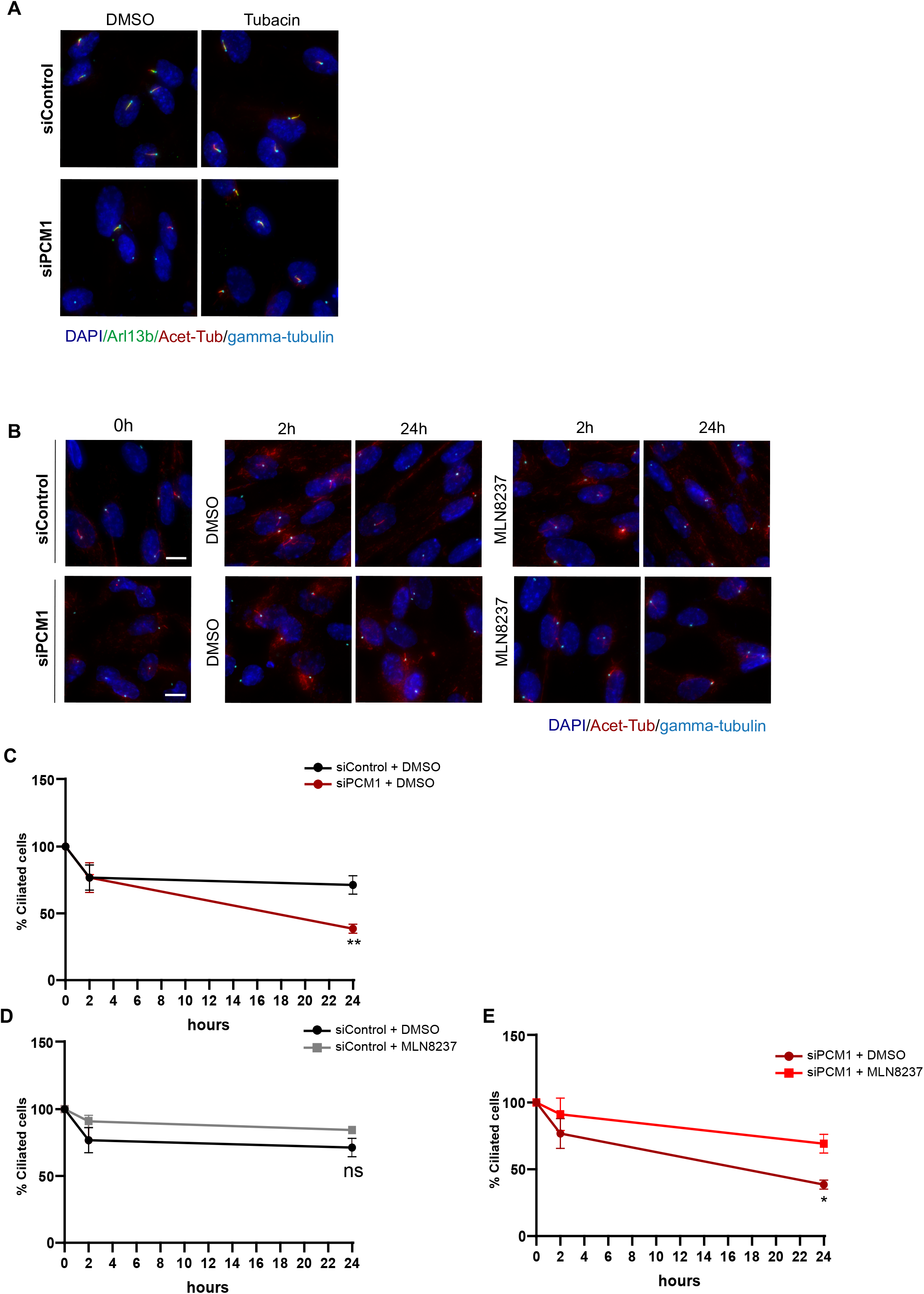
Analysis of cilium assembly and disassembly in control and PCM1-depleted cells treated with AURKA or HDAC6 inhibitor. **A.** Representative immunofluorescence images for cilium assembly experiments. RPE1 C=cells were transfected with control or PCM1 siRNA for 48 h and treated with DMSO (vehicle control) or 2 μM tubacin in serum starvation medium for 24 h. Cells were fixed and immunostained with acetylated tubulin antibody to mark the primary cilium and gamma tubulin antibody to mark the centrosome. DNA was stained with DAPI. Scale bar, 10 μm **B.** Representative immunofluorescence images for cilium disassembly experiments. RPE1 cells were transfected with control or PCM1 siRNA for 48 h, serum starved for 24 h and treated with DMSO (vehicle control) or 0.5 μM MLN8237 in serum stimulation medium for 2 h and 24 h. Cells were fixed and immunostained with acetylated tubulin antibody to mark the primary cilium and gamma tubulin antibody to mark the centrosome. DNA was stained with DAPI. Scale bar, 10 μm **C.** Graph represents the quantification of cilium disassembly in control and PCM1 depleted and DMSO treated cells. X-axis indicates the hours after serum stimulation. Data represent mean value from three experiments per condition ± SEM (**p<0.01) **D.** Graph represents the quantification of cilium disassembly in control depleted cells treated with DMSO or 1 μM MLN8237. X-axis indicates the hours after serum stimulation. Data represent mean value from three experiments per condition ± SEM (ns: non-significant) **E.** Graph represents the quantification of cilium disassembly in PCM1 depleted cells treated with DMSO or 1 μM MLN8237. X-axis indicates the hours after serum stimulation. Data represent mean value from three experiments per condition ± SEM (*p<0.05)

## References

Arslanhan, M. D., Gulensoy, D. & Firat-Karalar, E. N. 2020. A Proximity Mapping Journey into the Biology of the Mammalian Centrosome/Cilium Complex. Cells, 9.

Aydin, O. Z., Taflan, S. O., Gurkaslar, C. & Firat-Karalar, E. N. 2020. Acute inhibition of centriolar satellite function and positioning reveals their functions at the primary cilium. PLoS Biol, 18, e3000679.

Barr, A. R. & Gergely, F. 2007. Aurora-A: the maker and breaker of spindle poles. J Cell Sci, 120, 2987–96.

Bayliss, R., Sardon, T., Vernos, I. & Conti, E. 2003. Structural basis of Aurora-A activation by TPX2 at the mitotic spindle. Mol Cell, 12, 851–62.

Bertolin, G., Bulteau, A. L., Alves-Guerra, M. C., Burel, A., Lavault, M. T., Gavard, O., Le Bras, S., Gagne, J. P., Poirier, G. G., Le Borgne, R., Prigent, C. & Tramier, M. 2018. Aurora kinase A localises to mitochondria to control organelle dynamics and energy production. Elife, 7.

Bertolin, G. & Tramier, M. 2020. Insights into the non-mitotic functions of Aurora kinase A: more than just cell division. Cell Mol Life Sci, 77, 1031–1047.

Bhogaraju, S., Cajanek, L., Fort, C., Blisnick, T., Weber, K., Taschner, M., Mizuno, N., Lamla, S., Bastin, P., Nigg, E. A. & Lorentzen, E. 2013. Molecular basis of tubulin transport within the cilium by IFT74 and IFT81. Science, 341, 1009–12.

Buchel, G., Carstensen, A., Mak, K. Y., Roeschert, I., Leen, E., Sumara, O., Hofstetter, J., Herold, S., Kalb, J., Baluapuri, A., Poon, E., Kwok, C., Chesler, L., Maric, H. M., Rickman, D. S., Wolf, E., Bayliss, R., Walz, S. & Eilers, M. 2017. Association with Aurora-A Controls N-MYC-Dependent Promoter Escape and Pause Release of RNA Polymerase II during the Cell Cycle. Cell Rep, 21, 3483–3497.

Byrum, A. K., Carvajal-Maldonado, D., Mudge, M. C., Valle-Garcia, D., Majid, M. C., Patel, R., Sowa, M. E., Gygi, S. P., Harper, J. W., Shi, Y., Vindigni, A. & Mosammaparast, N. 2019. Mitotic regulators TPX2 and Aurora A protect DNA forks during replication stress by counteracting 53BP1 function. J Cell Biol, 218, 422–432.

Cheeseman, I. M., Anderson, S., Jwa, M., Green, E. M., Kang, J., Yates, J. R., 3RD, Chan, C. S., Drubin, D. G. & Barnes, G. 2002. Phospho-regulation of kinetochore-microtubule attachments by the Aurora kinase Ipl1p. Cell, 111, 163–72.

Chen, S. H. & Lahav, G. 2016. Two is better than one; toward a rational design of combinatorial therapy. Curr Opin Struct Biol, 41, 145–150.

Conkar, D., Bayraktar, H. & Firat-Karalar, E. N. 2019. Centrosomal and ciliary targeting of CCDC66 requires cooperative action of centriolar satellites, microtubules and molecular motors. Sci Rep, 9, 14250.

Conkar, D. & Firat-Karalar, E. N. 2020. Microtubule-associated proteins and emerging links to primary cilium structure, assembly, maintenance, and disassembly. FEBS J.

Deretic, J., Kerr, A. & Welburn, J. P. I. 2019. A rapid computational approach identifies SPICE1 as an Aurora kinase substrate. Mol Biol Cell, 30, 312–323.

Devaul, N., Koloustroubis, K., Wang, R. & Sperry, A. O. 2017. A novel interaction between kinase activities in regulation of cilia formation. BMC Cell Biol, 18, 33.

Dutertre, S., Cazales, M., Quaranta, M., Froment, C., Trabut, V., Dozier, C., Mirey, G., Bouche, J. P., Theis-Febvre, N., Schmitt, E., Monsarrat, B., Prigent, C. & Ducommun, B. 2004. Phosphorylation of CDC25B by Aurora-A at the centrosome contributes to the G2-M transition. J Cell Sci, 117, 2523–31.

Firat-Karalar, E. N., Rauniyar, N., Yates, J. R., 3RD & Stearns, T. 2014. Proximity interactions among centrosome components identify regulators of centriole duplication. Curr Biol, 24, 664–70.

Gheiratmand, L., Coyaud, E., Gupta, G. D., Laurent, E. M., Hasegan, M., Prosser, S. L., Goncalves, J., Raught, B. & Pelletier, L. 2019. Spatial and proteomic profiling reveals centrosome-independent features of centriolar satellites. EMBO J.

Gingras, A. C., Abe, K. T. & Raught, B. 2019. Getting to know the neighborhood: using proximity-dependent biotinylation to characterize protein complexes and map organelles. Curr Opin Chem Biol, 48, 44–54.

Glover, D. M., Leibowitz, M. H., Mclean, D. A. & Parry, H. 1995. Mutations in aurora prevent centrosome separation leading to the formation of monopolar spindles. Cell, 81, 95–105.

Golemis, E. A., Scheet, P., Beck, T. N., Scolnick, E. M., Hunter, D. J., Hawk, E. & Hopkins, N. 2018. Molecular mechanisms of the preventable causes of cancer in the United States. Genes Dev, 32, 868–902.

Gupta, G. D., Coyaud, E., Goncalves, J., Mojarad, B. A., Liu, Y., Wu, Q., Gheiratmand, L., Comartin, D., Tkach, J. M., Cheung, S. W., Bashkurov, M., Hasegan, M., Knight, J. D., Lin, Z. Y., Schueler, M., Hildebrandt, F., Moffat, J., Gingras, A. C., Raught, B. & Pelletier, L. 2015. A Dynamic Protein Interaction Landscape of the Human Centrosome-Cilium Interface. Cell, 163, 1484–99.

Gurkaslar, H. K., Culfa, E., Arslanhan, M. D., Lince-Faria, M. & Firat-Karalar, E. N. 2020. CCDC57 Cooperates with Microtubules and Microcephaly Protein CEP63 and Regulates Centriole Duplication and Mitotic Progression. Cell Rep, 31, 107630.

Hasanov, E., Chen, G., Chowdhury, P., Weldon, J., Ding, Z., Jonasch, E., Sen, S., Walker, C. L. & Dere, R. 2017. Ubiquitination and regulation of AURKA identifies a hypoxia-independent E3 ligase activity of VHL. Oncogene, 36, 3450–3463.

Higgins, M., Obaidi, I. & Mcmorrow, T. 2019. Primary cilia and their role in cancer. Oncol Lett, 17, 3041–3047.

Hirota, T., Kunitoku, N., Sasayama, T., Marumoto, T., Zhang, D., Nitta, M., Hatakeyama, K. & Saya, H. 2003. Aurora-A and an interacting activator, the LIM protein Ajuba, are required for mitotic commitment in human cells. Cell, 114, 585–98.

Hornbeck, P. V., Zhang, B., Murray, B., Kornhauser, J. M., Latham, V. & Skrzypek, E. 2015. PhosphoSitePlus, 2014: mutations, PTMs and recalibrations. Nucleic Acids Res, 43, D512–20.

Hutterer, A., Berdnik, D., Wirtz-Peitz, F., Zigman, M., Schleiffer, A. & Knoblich, J. A. 2006. Mitotic activation of the kinase Aurora-A requires its binding partner Bora. Dev Cell, 11, 147–57.

Inaba, H., Goto, H., Kasahara, K., Kumamoto, K., Yonemura, S., Inoko, A., Yamano, S., Wanibuchi, H., He, D., Goshima, N., Kiyono, T., Hirotsune, S. & Inagaki, M. 2016. Ndel1 suppresses ciliogenesis in proliferating cells by regulating the trichoplein-Aurora A pathway. J Cell Biol, 212, 409–23.

Inoko, A., Matsuyama, M., Goto, H., Ohmuro-Matsuyama, Y., Hayashi, Y., Enomoto, M., Ibi, M., Urano, T., Yonemura, S., Kiyono, T., Izawa, I. & Inagaki, M. 2012. Trichoplein and Aurora A block aberrant primary cilia assembly in proliferating cells. J Cell Biol, 197, 391–405.

Jackson, P. K. 2011. Do cilia put brakes on the cell cycle? Nat Cell Biol, 13, 340–2.

Joachim, J., Razi, M., Judith, D., Wirth, M., Calamita, E., Encheva, V., Dynlacht, B. D., Snijders, A. P., O’reilly, N., Jefferies, H. B. J. & Tooze, S. A. 2017. Centriolar Satellites Control GABARAP Ubiquitination and GABARAP-Mediated Autophagy. Curr Biol, 27, 2123–2136 e7.

Joukov, V. & De Nicolo, A. 2018. Aurora-PLK1 cascades as key signaling modules in the regulation of mitosis. Sci Signal, 11.

Kashatus, D. F., Lim, K. H., Brady, D. C., Pershing, N. L., Cox, A. D. & Counter, C. M. 2011. RALA and RALBP1 regulate mitochondrial fission at mitosis. Nat Cell Biol, 13, 1108–15.

Kettenbach, A. N., Schweppe, D. K., Faherty, B. K., Pechenick, D., Pletnev, A. A. & Gerber, S. A. 2011. Quantitative phosphoproteomics identifies substrates and functional modules of Aurora and Polo-like kinase activities in mitotic cells. Sci Signal, 4, rs5.

Kim, J., Krishnaswami, S. R. & Gleeson, J. G. 2008. CEP290 interacts with the centriolar satellite component PCM-1 and is required for Rab8 localization to the primary cilium. Hum Mol Genet, 17, 3796–805.

Kim, S., Zaghloul, N. A., Bubenshchikova, E., Oh, E. C., Rankin, S., Katsanis, N., Obara, T. & Tsiokas, L. 2011. Nde1-mediated inhibition of ciliogenesis affects cell cycle re-entry. Nat Cell Biol, 13, 351–60.

Kinzel, D., Boldt, K., Davis, E. E., Burtscher, I., Trumbach, D., Diplas, B., Attie-Bitach, T., Wurst, W., Katsanis, N., Ueffing, M. & Lickert, H. 2010. Pitchfork regulates primary cilia disassembly and left-right asymmetry. Dev Cell, 19, 66–77.

Kiseleva, A. A., Korobeynikov, V. A., Nikonova, A. S., Zhang, P., Makhov, P., Deneka, A. Y., Einarson, M. B., Serebriiskii, I. G., Liu, H., Peterson, J. R. & Golemis, E. A. 2019. Unexpected Activities in Regulating Ciliation Contribute to Off-target Effects of Targeted Drugs. Clin Cancer Res, 25, 4179–4193.

Korobeynikov, V., Deneka, A. Y. & Golemis, E. A. 2017. Mechanisms for nonmitotic activation of Aurora-A at cilia. Biochem Soc Trans, 45, 37–49.

Kufer, T. A., Sillje, H. H., Korner, R., Gruss, O. J., Meraldi, P. & Nigg, E. A. 2002. Human TPX2 is required for targeting Aurora-A kinase to the spindle. J Cell Biol, 158, 617–23.

Lambert, J. P., Tucholska, M., Go, C., Knight, J. D. & Gingras, A. C. 2014. Proximity biotinylation and affinity purification are complementary approaches for the interactome mapping of chromatin-associated protein complexes. J Proteomics.

Leroy, P. J., Hunter, J. J., Hoar, K. M., Burke, K. E., Shinde, V., Ruan, J., Bowman, D., Galvin, K. & Ecsedy, J. A. 2007. Localization of human TACC3 to mitotic spindles is mediated by phosphorylation on Ser558 by Aurora A: a novel pharmacodynamic method for measuring Aurora A activity. Cancer Res, 67, 5362–70.

Levinson, N. M. 2018. The multifaceted allosteric regulation of Aurora kinase A. Biochem J, 475, 2025–2042.

Li, A., Saito, M., Chuang, J. Z., Tseng, Y. Y., Dedesma, C., Tomizawa, K., Kaitsuka, T. & Sung, C. H. 2011. Ciliary transition zone activation of phosphorylated Tctex-1 controls ciliary resorption, S-phase entry and fate of neural progenitors. Nat Cell Biol, 13, 402–11.

Liem, K. F., JR., He, M., Ocbina, P. J. & Anderson, K. V. 2009. Mouse Kif7/Costal2 is a cilia-associated protein that regulates Sonic hedgehog signaling. Proc Natl Acad Sci U S A, 106, 13377–82.

Liu, X., Salokas, K., Tamene, F., Jiu, Y., Weldatsadik, R. G., Ohman, T. & Varjosalo, M. 2018. An AP-MS- and BioID-compatible MAC-tag enables comprehensive mapping of protein interactions and subcellular localizations. Nat Commun, 9, 1188.

Mellacheruvu, D., Wright, Z., Couzens, A. L., Lambert, J. P., St-Denis, N. A., Li, T., Miteva, Y. V., Hauri, S., Sardiu, M. E., Low, T. Y., Halim, V. A., Bagshaw, R. D., Hubner, N. C., Al-Hakim, A., Bouchard, A., Faubert, D., Fermin, D., Dunham, W. H., Goudreault, M., Lin, Z. Y., Badillo, B. G., Pawson, T., Durocher, D., Coulombe, B., Aebersold, R., Superti-Furga, G., Colinge, J., Heck, A. J., Choi, H., Gstaiger, M., Mohammed, S., Cristea, I. M., Bennett, K. L., Washburn, M. P., Raught, B., Ewing, R. M., Gingras, A. C. & Nesvizhskii, A. I. 2013. The CRAPome: a contaminant repository for affinity purification-mass spectrometry data. Nat Methods, 10, 730–6.

Mirvis, M., Stearns, T. & James Nelson, W. 2018. Cilium structure, assembly, and disassembly regulated by the cytoskeleton. Biochem J, 475, 2329–2353.

Mori, D., Yano, Y., Toyo-Oka, K., Yoshida, N., Yamada, M., Muramatsu, M., Zhang, D., Saya, H., Toyoshima, Y. Y., Kinoshita, K., Wynshaw-Boris, A. & Hirotsune, S. 2007. NDEL1 phosphorylation by Aurora-A kinase is essential for centrosomal maturation, separation, and TACC3 recruitment. Mol Cell Biol, 27, 352–67.

Nepusz, T., Yu, H. & Paccanaro, A. 2012. Detecting overlapping protein complexes in protein-protein interaction networks. Nat Methods, 9, 471–2.

Nikonova, A. S., Astsaturov, I., Serebriiskii, I. G., Dunbrack, R. L., JR. & Golemis, E. A. 2013. Aurora A kinase (AURKA) in normal and pathological cell division. Cell Mol Life Sci, 70, 661–87.

Odabasi, E., Batman, U. & Firat-Karalar, E. N. 2020. Unraveling the mysteries of centriolar satellites: time to rewrite the textbooks about the centrosome/cilium complex. Mol Biol Cell, 31, 866–872.

Odabasi, E., Gul, S., Kavakli, I. H. & Firat-Karalar, E. N. 2019. Centriolar satellites are required for efficient ciliogenesis and ciliary content regulation. EMBO Rep.

Otto, T., Horn, S., Brockmann, M., Eilers, U., Schuttrumpf, L., Popov, N., Kenney, A. M., Schulte, J. H., Beijersbergen, R., Christiansen, H., Berwanger, B. & Eilers, M. 2009. Stabilization of N-Myc is a critical function of Aurora A in human neuroblastoma. Cancer Cell, 15, 67–78.

Pan, J., Wang, Q. & Snell, W. J. 2004. An aurora kinase is essential for flagellar disassembly in Chlamydomonas. Dev Cell, 6, 445–51.

Pejskova, P., Reilly, M. L., Bino, L., Bernatik, O., Dolanska, L., Ganji, R. S., Zdrahal, Z., Benmerah, A. & Cajanek, L. 2020. KIF14 controls ciliogenesis via regulation of Aurora A and is important for Hedgehog signaling. J Cell Biol, 219.

Plotnikova, O. V., Nikonova, A. S., Loskutov, Y. V., Kozyulina, P. Y., Pugacheva, E. N. & Golemis, E. A. 2012. Calmodulin activation of Aurora-A kinase (AURKA) is required during ciliary disassembly and in mitosis. Mol Biol Cell, 23, 2658–70.

Prosser, S. L. & Pelletier, L. 2020. Centriolar satellite biogenesis and function in vertebrate cells. J Cell Sci, 133.

Pugacheva, E. N., Jablonski, S. A., Hartman, T. R., Henske, E. P. & Golemis, E. A. 2007. HEF1-dependent Aurora A activation induces disassembly of the primary cilium. Cell, 129, 1351–63.

Reiter, J. F. & Leroux, M. R. 2017. Genes and molecular pathways underpinning ciliopathies. Nat Rev Mol Cell Biol, 18, 533–547.

Roux, K. J. 2013. Marked by association: techniques for proximity-dependent labeling of proteins in eukaryotic cells. Cell Mol Life Sci, 70, 3657–64.

Roux, K. J., Kim, D. I., Raida, M. & Burke, B. 2012. A promiscuous biotin ligase fusion protein identifies proximal and interacting proteins in mammalian cells. J Cell Biol, 196, 801–10.

Santamaria, A., Wang, B., Elowe, S., Malik, R., Zhang, F., Bauer, M., Schmidt, A., Sillje, H. H., Korner, R. & Nigg, E. A. 2011. The Plk1-dependent phosphoproteome of the early mitotic spindle. Mol Cell Proteomics, 10, M110004457.

Schneider, C. A., Rasband, W. S. & Eliceiri, K. W. 2012. NIH Image to ImageJ: 25 years of image analysis. Nat Methods, 9, 671–5.

Seiler, C. Y., Park, J. G., Sharma, A., Hunter, P., Surapaneni, P., Sedillo, C., Field, J., Algar, R., Price, A., Steel, J., Throop, A., Fiacco, M. & Labaer, J. 2014. DNASU plasmid and PSI:Biology-Materials repositories: resources to accelerate biological research. Nucleic Acids Res, 42, D1253–60.

Spektor, A., Tsang, W. Y., Khoo, D. & Dynlacht, B. D. 2007. Cep97 and CP110 suppress a cilia assembly program. Cell, 130, 678–90.

Stark, C., Breitkreutz, B. J., Reguly, T., Boucher, L., Breitkreutz, A. & Tyers, M. 2006. BioGRID: a general repository for interaction datasets. Nucleic Acids Res, 34, D535–9.

Stowe, T. R., Wilkinson, C. J., Iqbal, A. & Stearns, T. 2012. The centriolar satellite proteins Cep72 and Cep290 interact and are required for recruitment of BBS proteins to the cilium. Mol Biol Cell, 23, 3322–35.

Tang, A., Gao, K., Chu, L., Zhang, R., Yang, J. & Zheng, J. 2017. Aurora kinases: novel therapy targets in cancers. Oncotarget, 8, 23937–23954.

Treekitkarnmongkol, W., Katayama, H., Kai, K., Sasai, K., Jones, J. C., Wang, J., Shen, L., Sahin, A. A., Gagea, M., Ueno, N. T., Creighton, C. J. & Sen, S. 2016. Aurora kinase-A overexpression in mouse mammary epithelium induces mammary adenocarcinomas harboring genetic alterations shared with human breast cancer. Carcinogenesis, 37, 1180–1189.

Van De Mark, D., Kong, D., Loncarek, J. & Stearns, T. 2015. MDM1 is a microtubule-binding protein that negatively regulates centriole duplication. Mol Biol Cell, 26, 3788–802.

Walter, A. O., Seghezzi, W., Korver, W., Sheung, J. & Lees, E. 2000. The mitotic serine/threonine kinase Aurora2/AIK is regulated by phosphorylation and degradation. Oncogene, 19, 4906–16.

Wang, G., Chen, Q., Zhang, X., Zhang, B., Zhuo, X., Liu, J., Jiang, Q. & Zhang, C. 2013a. PCM1 recruits Plk1 to the pericentriolar matrix to promote primary cilia disassembly before mitotic entry. J Cell Sci, 126, 1355–65.

Wang, L., Lee, K., Malonis, R., Sanchez, I. & Dynlacht, B. D. 2016. Tethering of an E3 ligase by PCM1 regulates the abundance of centrosomal KIAA0586/Talpid3 and promotes ciliogenesis. Elife, 5.

Wang, W. J., Tay, H. G., Soni, R., Perumal, G. S., Goll, M. G., Macaluso, F. P., Asara, J. M., Amack, J. D. & Tsou, M. F. 2013b. CEP162 is an axoneme-recognition protein promoting ciliary transition zone assembly at the cilia base. Nat Cell Biol, 15, 591–601.

Wang, X., Zhou, Y. X., Qiao, W., Tominaga, Y., Ouchi, M., Ouchi, T. & Deng, C. X. 2006. Overexpression of aurora kinase A in mouse mammary epithelium induces genetic instability preceding mammary tumor formation. Oncogene, 25, 7148–58.

Willems, E., Dedobbeleer, M., Digregorio, M., Lombard, A., Lumapat, P. N. & Rogister, B. 2018. The functional diversity of Aurora kinases: a comprehensive review. Cell Div, 13, 7.

Zhang, D., Hirota, T., Marumoto, T., Shimizu, M., Kunitoku, N., Sasayama, T., Arima, Y., Feng, L., Suzuki, M., Takeya, M. & Saya, H. 2004. Cre-loxP-controlled periodic Aurora-A overexpression induces mitotic abnormalities and hyperplasia in mammary glands of mouse models. Oncogene, 23, 8720–30.

Zhang, D., Shimizu, T., Araki, N., Hirota, T., Yoshie, M., Ogawa, K., Nakagata, N., Takeya, M. & Saya, H. 2008. Aurora A overexpression induces cellular senescence in mammary gland hyperplastic tumors developed in p53-deficient mice. Oncogene, 27, 4305–14.

Zybailov, B., Mosley, A. L., Sardiu, M. E., Coleman, M. K., Florens, L. & Washburn, M. P. 2006. Statistical analysis of membrane proteome expression changes in Saccharomyces cerevisiae. J Proteome Res, 5, 2339–47.

